# Integrative Chemical Genetics Platform Identifies Condensate Modulators Linked to Neurological Disorders

**DOI:** 10.1101/2025.06.07.658469

**Authors:** Dylan Poch, Chandrayee Mukherjee, Sunanda Mallik, Vanessa Todorow, E.F. Elsiena Kuiper, Nalini Dhingra, Yulia V Surovtseva, Christian Schlieker

## Abstract

Aberrant biomolecular condensates are implicated in neurological disorders including ALS, frontotemporal dementia, and DYT1 dystonia, yet approaches to systematically identify their modulators remain limited. Here we establish MLF2 as a versatile condensate biomarker and develop CondenScreen, an integrated high-content screening and bioinformatic pipeline enabling identification of condensate modulators across chemical and genetic space. Screening 1,760 FDA-approved compounds in a cellular DYT1 dystonia model, we identify drugs that alter aberrant condensate properties, validating the platform for condensate-targeted drug discovery. In parallel, a genome-wide CRISPR/Cas9 screen links condensate accumulation to microcephaly genes and more than 10 additional neurodevelopmental disorders. Machine learning and confocal imaging resolves distinct condensate phenotypes: loss of microcephaly-associated ZNF335 results in nucleoplasmic condensates, whereas RNF26 deletion produces nuclear envelope condensates that phenocopy hallmarks of torsin deficiency. Our study provides a scalable resource for identifying corrective modulators of condensates and establishes a link between nuclear condensate accumulation and neurodevelopmental disorders.

## Introduction

Dysregulation of physiological condensates is a driver for many neurological disorders, including Amyotrophic Lateral Sclerosis (ALS), Frontotemporal Dementia (FTD), and DYT1 dystonia.^1–5^ Biomolecular condensates, often made from a milieu of hundreds of different proteins, consist of a thermodynamically favorable separation between a dense and dilute phase.^6,7^ When properly regulated, biomolecular condensates allow efficient, fine-tuned microenvironments, ensuring maintenance of cellular homeostasis while buffering against external perturbations.^8–10^ However, in pathological states referred to as condensatopathies, these condensates exhibit reduced dynamicity, leading to protein aggregation or amyloid formation that is not easily reversible.^11–13^ It is estimated that up to one-third of common diseases can be linked to condensate-forming proteins.^14^ Determining mechanisms and cellular activities of these processes is essential for devising therapeutic strategies that tune condensate properties to counteract pathological transitions.

One favorable phase-state is observed in the central channel of the nuclear pore complex (NPC). The NPC serves as the intermediary gate at which nuclear transcription is separated from cytoplasmic translation in eukaryotic organisms. Within the central channel of the NPC, phenylalanine-glycine nucleoporins (FG-nups) have a high proportion of intrinsically disordered regions that set up a selective phase, contributing to the selective barrier of the NPC.^15–18^ Disruptions to NPC assembly or nucleocytoplasmic transport have been strongly implicated in neurological diseases, including DYT1 dystonia,^1,19,20^ a debilitating movement disorder characterized by involuntary muscle contractions and painful sustained postures.^21,22^

DYT1 dystonia is caused by a mutation within the TOR1A gene that disrupts activation of the encoded AAA+ ATPase,^23–27^ TorsinA, resulting in the formation of pathological nuclear envelope condensates.^1,19,20,28,29^ As with many condensatopathies, there are only palliative interventions, with no curative or preventative treatments for dystonia.^21,22^ These nuclear envelope condensates accumulate K48-linked polyubiquitinylated proteins that evade proteasomal degradation and sequester chaperone proteins, causing cellular proteotoxicity.^1,20,30^ Thus, regulating aberrant nuclear condensates by either direct dissolution or modulation of its proteotoxic properties represents a practical therapeutic approach. However, current limitations to high-throughput screening campaigns targeting condensatopathies include a lack of well-defined condensate targets and broadly applicable condensate biomarkers.

In this study, we establish Myeloid leukemia factor 2 (MLF2) as a robust reporter that is enriched in proteotoxic nuclear condensates and stress granules, as evidenced by co-localization with TDP-43 and G3BP1. Using an MLF2 readout, we integrated complementary chemical and genetic high-content screening strategies to define the landscape underlying atypical nuclear condensates. First, a target-agonistic chemical screen in a cell-based DYT1 dystonia model identified several FDA-approved small molecules that modulate proteotoxic nuclear condensates, providing a tractable methodological framework for identifying therapeutics for condensatopathies. Second, a CRISPR/Cas9 knockout (KO) screen in wild-type cells, combined with deep learning approaches, identified RNF26 and ZNF335 as protective factors against distinct condensate phenotypes. Notably, 15 out of the top 50 genetic hits are associated with neurodevelopmental disorders, including seven directly linked to primary microcephaly or related congenital brain malformations. Together, these orthogonal approaches converge on a shared connection between nuclear condensate accumulation, perturbed nuclear proteostasis, and neurodevelopmental disorders, while establishing a scalable platform for therapeutic discovery.

## Results

### MLF2 is a Biomarker for Condensates Associated with ALS/FTD and DYT1 Dystonia

Having established the connection between impaired protein quality control and condensates in a torsin-deficient DYT1 dystonia HeLa cell model (4TorKO cells) (Figure 1A),^1,20^ we aimed to develop biomarkers to facilitate the identification of condensate modulators across other condensatopathies. MLF2 represents a major protein constituent of nuclear envelope condensates within 4TorKO cells^20^ and readily immerses into FG-nup condensates *in vitro*,^1^ likely due to its highly enriched Methionine and Arginine residues. Since many condensates share aromatic side chains as part of their molecular grammar,^31^ we previously suggested that MLF2 might localize to additional aberrant condensates through π-cation interactions as MLF2 is highly enriched in Arg residues that cannot be functionally replaced by charge-conserving Lys residues.^1^ Here, we have exploited these properties to visualize proteotoxic condensates by designing doxycycline (Dox) inducible fluorophore-derivatized variants of MLF2 as condensate biomarkers.

**Fig. 1.**
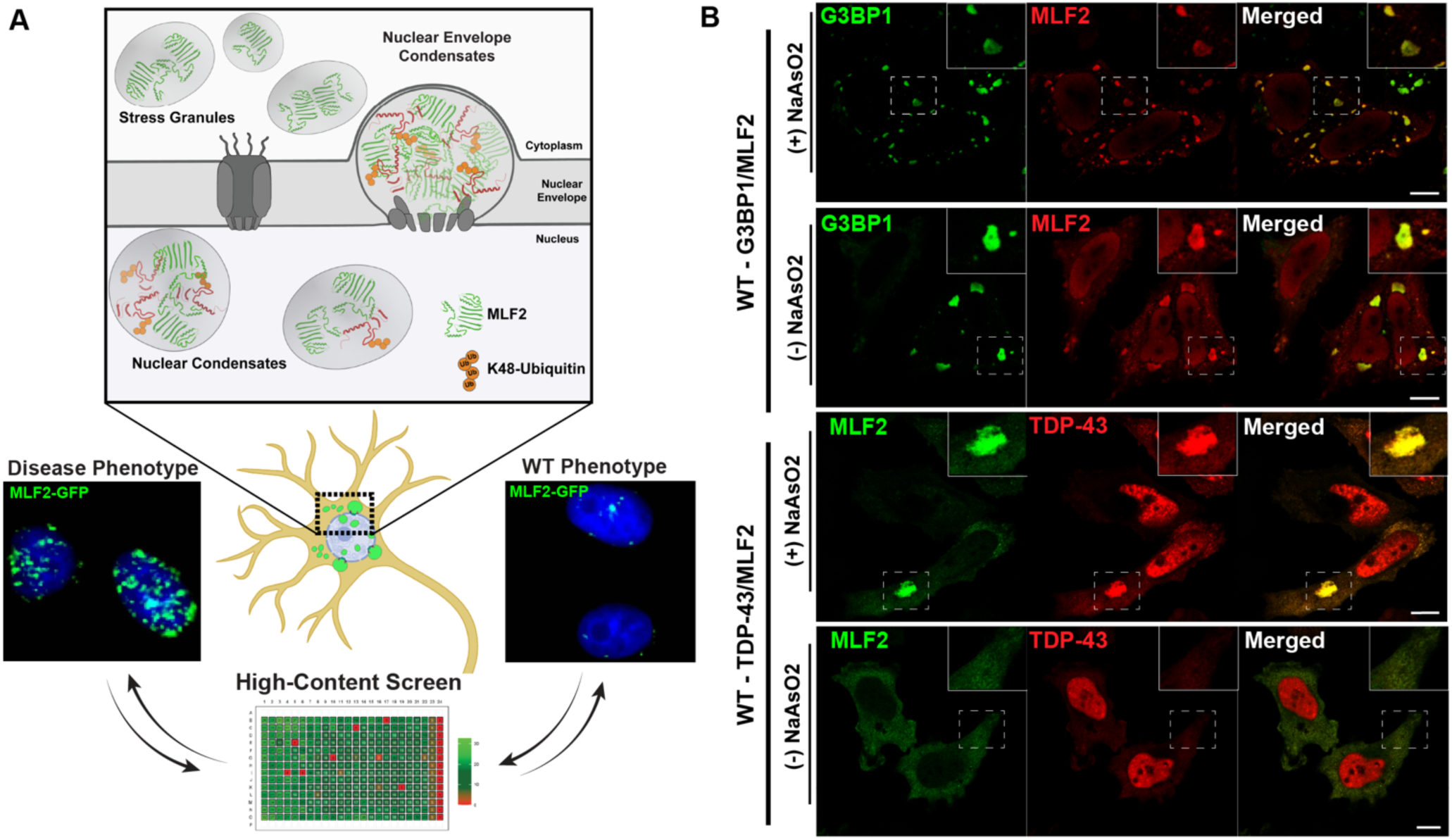
MLF2 is a biomarker for aberrant condensates across neurological disorders. (**A**) MLF2 detects both nuclear condensates and stress granules. Nuclear envelope condensates that sequester MLF2, FG-nups (not shown for clarity), and K48-Ubiquitin-enriched proteins have been identified in DYT1 dystonia models. The ability of MLF2 to immerse into condensates can be leveraged for high-content imaging platforms to systematically profile additional disease-relevant condensates. (**B**) MLF2 co-localizes with G3BP1 (top panels) and TDP-43 (bottom panels) after stress granule formation in response to oxidative stress (0.5 mM NaAsO_2_, 30 min).

Dysregulation of stress granules (SGs) has been pathologically associated with several neurological disorders, including ALS and FTD.^11^ To assess MLF2 localization to stress granules, we co-expressed the SG markers G3BP1 and TDP-43 in wild-type (WT) HeLa cells exposed to 0.5 mM sodium arsenite (NaAsO_2_) for 30 minutes, an established condition for SG induction.^32–35^ Upon NaAsO_2_ exposure, MLF2 exhibited strong co-localization with both G3BP1-GFP and TDP-43-tdTomato (Figure 1B). These findings suggest that MLF2 serves as a robust reporter of condensates associated with distinct neurological disorders.

### Establishing MLF2-GFP as a Condensate Reporter for High-Content Screening

Leveraging the condensate biomarker properties of MLF2, we developed a miniaturized, high-content assay using Dox-inducible MLF2-GFP cells in a 384-well format. This facilitated the design of phenotypic, high-content arrayed screens to identify chemical and genetic modulators of nuclear condensates. Specifically, each well contained either a small molecule applied to 4TorKO cells or synthetic single guide RNAs (sgRNAs) delivered to Cas9-integrated WT cells, designed to suppress or promote nuclear condensates, respectively. To analyze the high-resolution images (over 10k images/plate), we developed a computational condensate screening suite, *CondenScreen*. This CellProfiler and R script was designed for the rapid quantification and statistical analysis of condensates (Figure S1A).

While many robust computational resources are available for monitoring condensates (see *PhaseMetrics* and *Punctatools*)^36,37^, *CondenScreen* is specifically designed for large-scale, high-content screening analysis using a customizable *CellProfiler 4* ^38,39^ pipeline and *R* analysis script. To this end, *CondenScreen* quantifies condensates at both single-cell and population levels and provides a graphical user interface (GUI) allowing users to select plate size and layout, as well as tailoring downstream calculations for a signal “ON” vs signal “OFF” screening campaign. In addition, built-in quality control features automatically assess screen robustness using established statistical metrics, including the Z’-Score, BZ-Score, signal-to-background (SB), normalized effect, and coefficient of variation (CV). In addition, the platform has optional filters to flag potential false positives arising from either spatial plate effects (Figure S1B-S1D) and filter candidate compounds with known false positive liabilities, including pan-assay interference compounds (PAINS) and additional structural alert categories (Figure S1E). Integration of the MLF2-GFP inducible system with our *CondenScreen* analysis pipeline ensures rapid identification of condensate modulators.

### High-Content Screen Identifies FDA-approved Drugs that Modulate Condensates

To identify chemical matter that can modulate aberrant nuclear envelope condensates, we employed our MLF2-GFP inducible 4TorKO DYT1 cell model. We screened 1760 drugs from the Pharmakon library, comprising 1360 US and 400 international drugs, at a 10 µM working concentration. To minimize false positives due to differential reporter expression, Dox (0.5 µg/ml) was added for four hours, followed by an additional 2 hours with compound incubation.

*CondenScreen* reported an average Z’-score of 0.75 indicating pronounced separation between the positive (0.1% DMSO-treated WT HeLa) and negative (0.1% DMSO-treated 4TorKO HeLa) controls.^40^ Additionally, the ratio between the negative and positive controls (SB) was 950, while the SD and CV of DMSO-treated cells were 1.9% and 8.3%, respectively, further confirming assay robustness.

To control for plate-specific column, row, and edge effects, we assigned each compound a BZ-score based on its ability to decrease the number of nuclear MLF2-GFP foci (Figure 2A, see methods). The distribution of the average number of MLF2-GFP foci per condition was near-normally distributed (Figures 2B, 2C). Using a cutoff BZ-score of three, we identified 20 putative hit compounds that significantly depleted MLF2-GFP condensate foci. Top compounds included plumbagin, sanguinarium chloride, thiram, piroctone olamine, penfluridol, bronopol, thimerosal, triclosan, disulfiram, celastrol, dihydrocelastryl diacetate, and pyrithione zinc (PZ). A full list of the screened drugs can be found in the supplementary materials (Table S1).

**Fig. 2.**
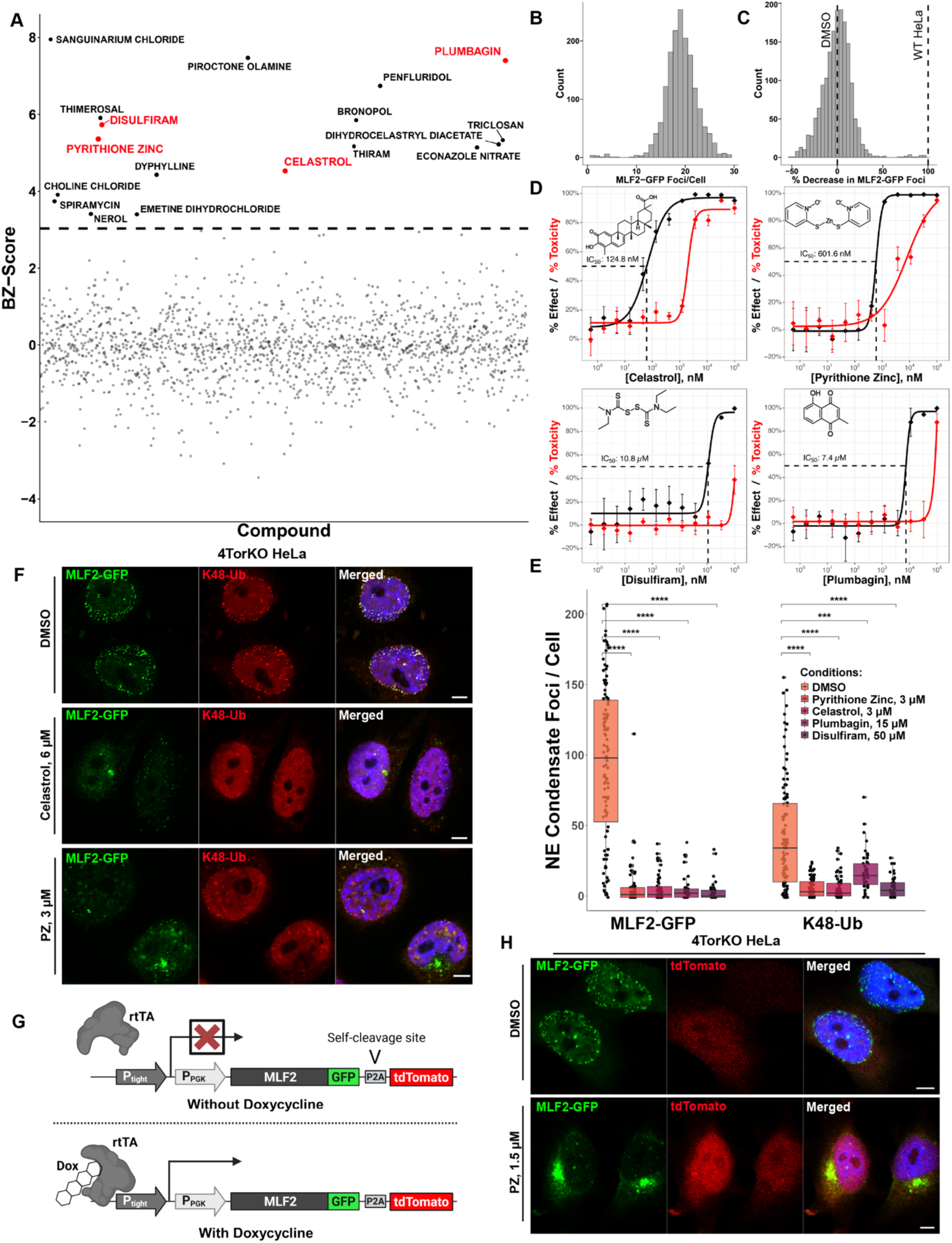
Chemical screen identifies drugs that modulate nuclear envelope condensates. (**A**) Drugs from Pharmakon 1760 screen decrease nuclear MLF2-GFP accumulation in 4TorKO HeLa cells. Compounds (circles) passing significance threshold (dotted-line, BZ-score > 3) are labeled purple, while red-labeled dots denote drugs selected for downstream validation. (**B**) Distribution in the average number of MLF2-GFP foci/cell and (**C**) percent decrease of MLF2-GFP foci per cell, normalized to the screening population. (**D**) Hit compounds are dose dependent. Determined IC_50_ from 12-point dose response (four orders of magnitude, n = 4). Black line represents % effect (decrease in MLF2-GFP foci), red line represents % toxicity relative to DMSO (four-parameter-log-logistic model). (**E**) Drugs modulate accumulation of K48-Ub proteins. Dots depict the number of MLF2-GFP (left, 6h Dox) or α-K48-ubiquitin (right) foci/cell after treatment with drug or DMSO. Asterisks indicate Bonferroni-corrected p-values calculated by Mann-Whitney U testing (***: p <= 0.001 ****: p <= 0.0001). (**F**) Representative immunofluorescence images of MLF2-GFP (green) and α-K48-ubiquitin (red) in torsin-deficient cells treated with DMSO compared to either PZ (3 µM) or celastrol (6 µM). (**G**) Schematic of inducible MLF2-GFP-P2A construct without (top) or with (bottom) Dox. (**H**) Confocal images of torsin-deficient cells expressing MLF2-GFP-P2A-tdTomato treated with either DMSO or 1.5 µM PZ treatments (6h). Scale bars, 5 µm.

Top-scoring compounds were resolubilized to construct a 12-point dose response (n = 4 replicates), covering over 5 orders of magnitude to determine the calculated IC_50_. (Figure 2D, Figure S2A). The normalized percent effect and percent toxicity (calculated as a function of viability, nuclear area, and nuclear morphology) were quantified for each concentration to determine potential therapeutic windows. Four structurally diverse drugs demonstrated a potent dose-dependent relationship, which included celastrol (IC_50_ = 124.8 nM), PZ (IC_50_ = 601.6 nM), plumbagin (IC_50_ = 7.4 µM), and disulfiram (IC_50_ = 10.8 µM) (Figure 2D).

### Pyrithione Zinc Disrupts K48-Ubiquitinylated Protein Accumulation within Nuclear Condensates

To determine the optimal concentration, we tested each of the four candidate compounds—celastrol, PZ, plumbagin, and disulfiram—at four tighter concentrations, including the IC_50_ itself, at 2x above/below the IC_50_, and 8x above the IC_50_ concentrations, using a 3 h compound treatment. The resulting boxplot reflects each compound’s effect at the single-cell level (Figure S2B). Consistent with the significant cell-cycle dependency of NE condensate formation previously demonstrated,^20^ we observed significant variability in the number of MLF2-GFP foci/cells. However, all compounds significantly reduced perinuclear MLF2-GFP accumulation at 2x and 8x their respective IC_50_ (Figure S2B).

To ensure consistent reduction across different condensate markers, we used K48-linked ubiquitin (K48) as an independent and endogenous biomarker based on our previous observation that 4TorKO cells have reduced protein turnover, resulting in the accumulation of polyubiquitinylated proteins within nuclear envelope condensates.^1,20^ Using immunofluorescence staining with α-K48 antibodies, all four compounds demonstrated a statistically significant decrease in endogenous, perinuclear K48 foci, replicating our results at reducing MLF2-GFP accumulation (Figure 2E). Between the optimal concentration profile of PZ and celastrol (Figure 2D), and the undesirable structural elements of plumbagin (quinone, Michael acceptor; Figure S1F), we continued to validate PZ and celastrol by assessing MLF2-GFP and endogenous K48 foci using confocal microscopy (Figure 2F). Importantly, this data validates that the mechanism of action either directly or indirectly mediates endogenous protein cargo within condensates.

A significant limitation to phenotype screening is the prevalence of identification of false positives that inhibit biomarker expression by non-specifically disrupting cellular protein transcription or translation. To circumvent this, we constructed a HeLa cell line with an inducible bicistronic MLF2-GFP-P2A-tdTomato reporter (Figure 2G). The P2A site induces ribosomal skipping, allowing the translation of two separate polypeptides, an N-terminal MLF2-GFP and a C-terminal tdTomato, both derived from the same mRNA transcript.^41^ We selected MLF2-GFP-P2A-tdTomato 4TorKO cells using Fluorescence-Activated Cell Sorting (FACS) (Figure S2C), before inducing MLF2-GFP expression and treating cells with either PZ (0.75 µM or 1.5 µM, 6 h) or celastrol (1.5 µM or 3 µM, 6 h). We quantified the number of MLF2-GFP foci and tdTomato expression between either uninduced cells treated with DMSO vehicle control, induced cells treated with DMSO, and induced cells treated with either 100 µg/mL cycloheximide, PZ, or celastrol. Concentrations of both celastrol and PZ at 1.5 µM had a significant decrease in the number of MLF2-GFP foci/cell compared with the DMSO control (Figure S2D). While PZ did not reduce the expression of secondary-tdTomato reporter (Figures 2H, S2E), celastrol slightly, albeit at a non-statistically significant level, decreased secondary reporter expression compared with DMSO-treated cells.

To address and quantify the potentially confounding effects of a compound affecting reporter abundance non-specifically, we developed an empirical metric, the Condensate Normalization Score (CNS). The CNS is based on the differential fluorescence between the condensate biomarker and secondary reporter. A high CNS-value indicates a reduction in condensate biomarkers with minimal impact on overall protein levels (see methods). Both celastrol- and PZ-treated cells showed a significantly increased percentage of cells with a median CNS exceeding 5-fold that of the DMSO control (Figure S2F), indicating a mechanism independent of reporter expression.

To assess the specificity of the top two compounds in modulating aberrant condensates, we examined their effects on nuclear Promyelocytic Leukemia (PML) bodies, physiological phase-separated compartments involved in protein quality control. As expected, treatment with 5% 1,6-hexanediol (5 minutes), a known disrupter of liquid-liquid phase-separated environments,^42^ significantly reduced PML body number (Figure S2H, S2G). In contrast, neither PZ nor celastrol reduced PML body number at a concentration of 1.5 µM that profoundly reduced MLF2 foci in our disease model (Figure S2H, S2G). Interestingly, treatment with PZ resulted in a statistically significant increase to the number of PML body puncta. Overall, this indicates that the activity of PZ and celastrol does not reflect nonspecific disruption of physiological nuclear condensates. However, visualization of condensates in 4TorKO cells using transmission electron microscopy (TEM) did not reveal full dissolution of nuclear envelope condensates after a 3 h treatment of 2 µM PZ (Figure S2I).

To identify the structural features responsible for the condensate-modifying activity of PZ, we examined whether this effect originates from its pyrithione moiety (2-Mercaptopyridine N-oxide) or the zinc ion. To this end, we treated 4TorKO cells with either DMSO vehicle control, PZ alone, or PZ with zinc chelator TPEN (N,N,Nʹ,Nʹ-*tetrakis*-(2-Pyridylmethyl)ethylenediamine) at a 3 molar equivalent.^43^ After zinc chelation with TPEN, the pyrithione compound failed to reduce MLF2-GFP condensate accumulation in 4TorKO cells (Figures S3A, S3B). Furthermore, we tested whether zinc alone is sufficient to deplete nuclear MLF2-GFP condensate foci. Due to its lower membrane permeability, 4TorKO cells were treated with higher concentrations of zinc sulfate monohydrate (ZnSO_4_). Treatment with 2 mM ZnSO_4_ alone decreased MLF2-GFP foci (Figure S3C, S3D) while maintaining tdTomato expression (Figure S3E), resulting in a higher relative CNS value (Figure S3F). Thus, PZ modulates nuclear condensates through a zinc-dependent mechanism.

### Genome-Wide CRISPR Screening Uncovers Condensate Modulators Implicated in Microcephaly and Neurodevelopmental Disorders

Having established a robust screening pipeline, we next performed a genome-wide CRISPR/Cas9 KO screen to identify genetic determinants of nuclear condensate accumulation. Knowledge of the corresponding activities may facilitate the identification of future pharmacological targets. We screened an arrayed library of 19,195 human genes using the Horizon Discovery Edit-R synthetic sgRNA collection. To increase KO efficiency, we designed and FACS-sorted a WT HeLa cell line that constitutively expresses Cas9 with Dox-inducible MLF2-GFP. Each well was reverse-transfected in a 384-well plate format using a pool of three sgRNAs targeting different regions of the same gene (Figure 3A). Cells were incubated for 72 h, and MLF2-GFP was induced by Dox 16 h before fixation (all within the 72 h window). After running all high-content images through our *CondenScreen* pipeline, the average Z’ value was 0.52, demonstrating a dynamic range between the positive (TOR1A/B sgRNAs) and negative (non-targeting (NT) sgRNA) controls. The calculated SB was 4.8, and the SD and CV of the NT control were 3.1 and 9.7, respectively. We determined the assay quality score to be sufficiently robust as indicated by similar scores in the literature.^34,44^

**Fig. 3.**
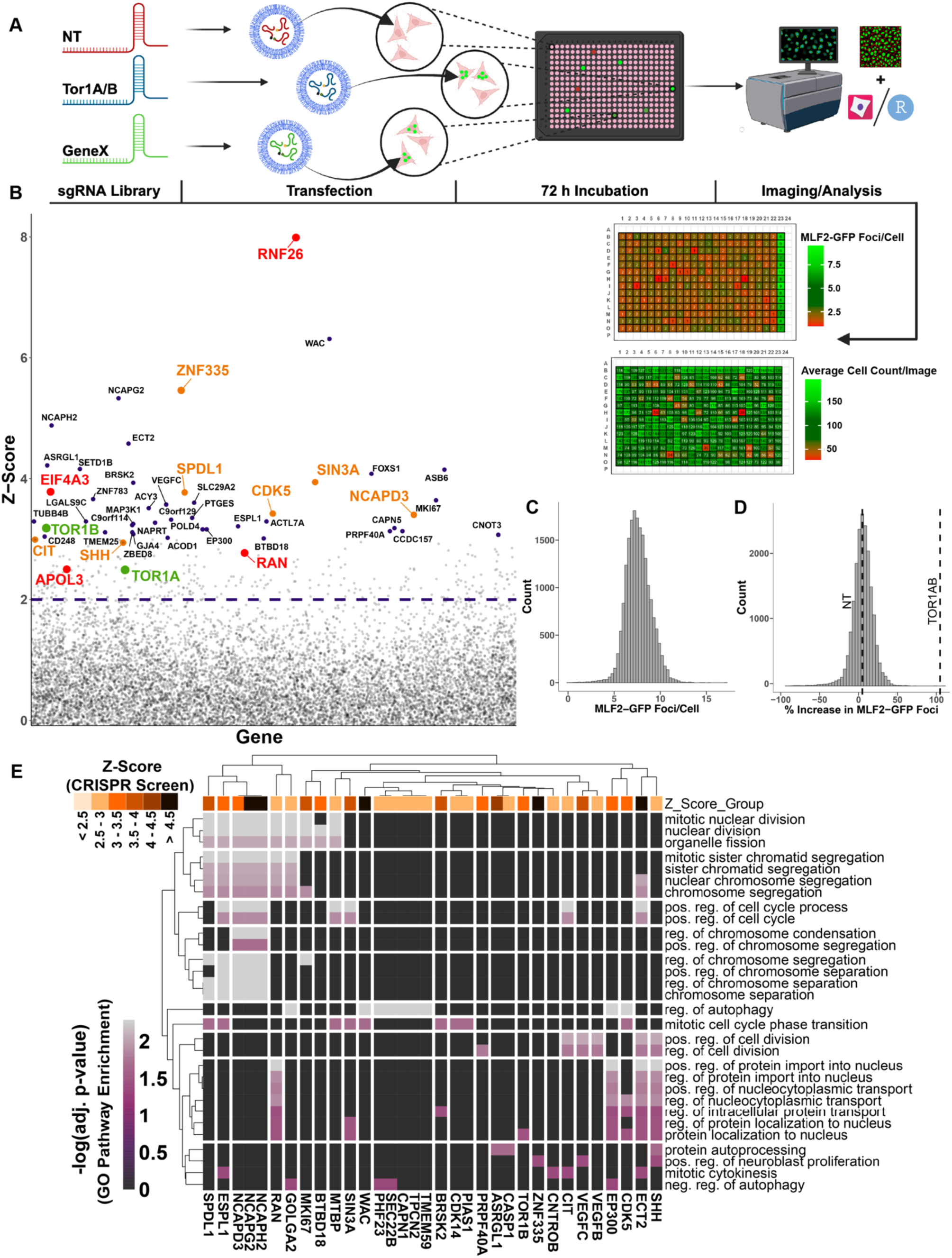
CRISPR KO Screen reveals genes that modulate nuclear condensates. (**A**) Schematic workflow of the genome-wide CRISPR KO screen using WT HeLa cells. Plates represent direct outputs provided by *CondenScreen*, indicating the number of MLF2-foci/cell (top plate) and cell count per image (bottom plate). (**B**) Genes hits from CRISPR KO screen associated with microcephaly modulate nuclear condensates. Dotted line represents significance threshold (Z-Score > 2) while highlighted and labeled genes have a Z-Score > 3. Green dots represent the TorsinA and TorsinB KO conditions; orange dots represent genes linked to microcephaly; and larger, red dots indicate genes of additional interest. (**C**) Distribution of the average number of MLF2-GFP foci/cell and (**D**) percent increase in MLF2-GFP foci/cell, normalized to the screening population. (**E**) Heatmap of the 30 highest scoring GO biological processes (rows) and associated genes (columns) from the top 100 candidate CRISPR KO hits. Vertical scale legend indicates −log(10) Benjamini-Hochberg adjusted p-value, with lighter regions representing higher significance. Scaled boxes (top) indicate the Z-Score of each gene from the CRISPR Screen, with darker colors indicating a higher Z-score (horizontal scale bar). Euclidean distance hierarchical clustering for GO-enriched biological pathways and genes is shown with the left and top dendrogram, respectively.

We defined a candidate gene as any gene deletion that resulted in a Z-score > 2 (Figure 3B). The top 10-ranked candidate genes were: RNF26, WAC, ZNF335, NCAPG2, NCAPH2, ECT2, ASRGL1, SETD1B, ASB6, and FOXS1 (see Table S2 for comprehensive list). Overall, the screen resulted in a near-normal distribution of the number of MLF2-GFP foci per cell (Figures 3C, 3D). Notably, we observed several genes linked to neurodevelopmental disorders, with high enrichment of microcephaly-associated genes (orange labeled dots, Figure 3B), according to human genetics annotated from OMIM^45^ and the NIH Genetic Testing Registry (GTR).^46^ Both homozygous and compound heterozygous mutations to one of our top hits, zinc finger protein 335 (ZNF335, also identified as MCPH10 or NIF1) have been documented to cause primary microcephaly.^47–54^ ZNF335 has been tied to the regulation of neurogenesis through modulation of the REST complex.^48^

Additional top hits of interest include, RNF26, an E3-ubiquitin ligase that has been implicated in regulating the endosomal system, serving as a “gatekeeper” for vesicles in the perinuclear ER cloud.^55,56^ Furthermore, RNF26 ubiquitinylates SQSTM1^55^ and has recently been tied to ER-sheet morphology and maintenance.^56^ Other hits include APOL3 (Z-Score = 2.5), a putative regulator of membrane dynamics with detergent-like bactericidal properties;^57,58^ Ran (Z-Score = 2.8), a GTP-binding nuclear protein; and EIF4A3/DDX48 (Z-Score 3.8), a DEAD-box ATPase. DEAD-box ATPases have recently been linked to the regulation of phase-separated membraneless organelles,^59^ indicating a potential role of EIF4A3 in modulating condensates. These condensate phenotypes observed across gene deletions highlight critical entry points into unexplored biology and promising directions for therapeutic discovery.

To determine if candidate genes are clustered into enriched genetic pathways, we hierarchically clustered the top 100 gene hits with their corresponding 30 enriched biological processes (q-value < 0.05) using Gene Ontology (GO) analysis^60^ (Figure 3E; interactive webpage available, see methods) and illustrated pathway interconnectedness (Figures S4A-S4C). We observed enrichment (10 out of 30 GO terms) of chromosomal and chromatid condensation, separation, or segregation pathways (Figures 3E, S4A). GO pathways were manually clustered into the following five categories: Autophagy, Cell Cycle, Chromosomal Regulation, Protein Transport, and Other (Figure S4B). Consistent with these enriched pathways, loss of protein homeostasis,^1,61^ a cell-cycle dependency of condensates,^20^ and disruption to nucleocytoplasmic transport^62,63^ have been documented in condensatopathies. Additionally, dysregulation of both cell cycle checkpoints and chromosomal regulation has been linked to primary microcephaly.^64–66^ Indeed, seven out of the top 50 gene hits have been linked to microcephaly.^45,46^ Beyond the enrichment of individual gene hits linked to microcephaly, encephalopathies, and other neurodevelopmental disorders, additional enriched pathways of interest included organelle fission and positive regulation of neuroblast proliferation.

To validate the three highest candidate genes, we knocked out ZNF335, RNF26, and the WW Domain Containing Adaptor with Coiled-Coil (WAC), and binned them according to the number of nuclear MLF2-GFP foci/cell (5-15, 15-30, or >30 foci/cell) (Figure S4D). While WAC depletion had only a modest effect, RNF26 KO and ZNF335 KO cells displayed pronounced nuclear condensate accumulation as indicated by MLF2-GFP accumulation (Figure S4D), with ZNF335 deficient cells resulting in the greatest number of condensates.

### Supervised Machine Learning Links Genetic Perturbations to Distinct Condensate Phenotypes

To resolve phenotypic distinctions and cluster genetic perturbations by their condensate morphology, potentially highlighting divergent biological roles, we developed a deep learning model to capture subtle morphological variations between condensates. Using a residual convolutional neural network (Figure S5A),^67^ we trained the model on single-cell patches derived from our strongest validated genetic hits to determine whether condensates differences could be computationally discriminated (Figure 4A). We employed a ResNet18 neural network backbone (ImageNet)^68^ to learn a regression mapping across three reference states: RNF26 KO (−1), non-targeting (0), and ZNF335 KO (+1). The model assigned each unlabeled cell a continuous phenotypic similarity score ranging from −1 (RNF26-like) to +1 (ZNF335-like)

**Fig. 4.**
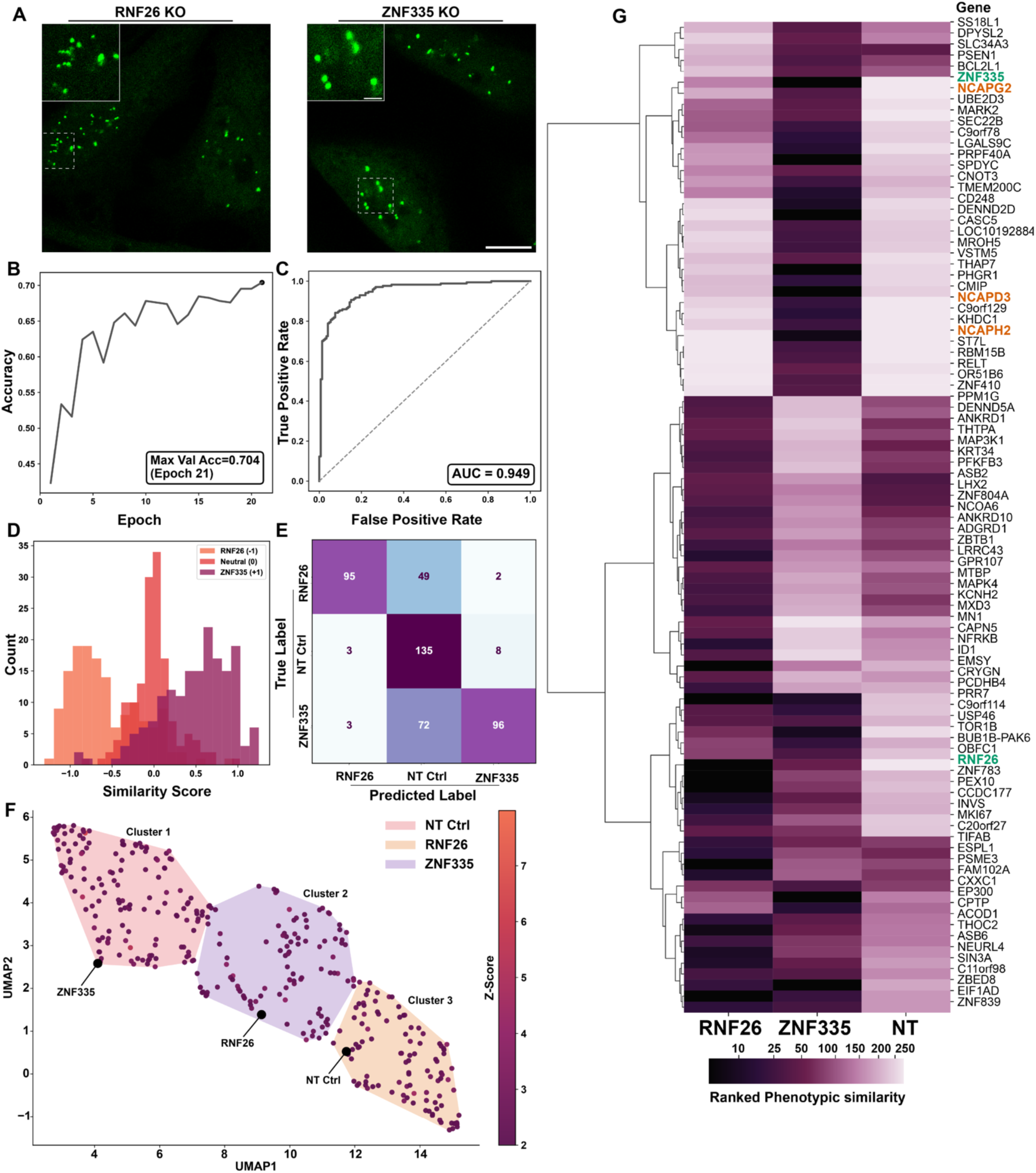
Deep learning model differentiates and clusters distinct condensate phenotypes. (**A**) RNF26 and ZNF335 have distinct MLF2-GFP condensate phenotypes. Representative confocal imaging of RNF26 KO (left) or ZNF335 KO (right) HeLa cells expressing MLF2-GFP (green). Scale bars, 10 µm. (**B**) Accuracy curve of model during training. Accuracy reflects the percentage of validation cells correctly classified as RNF26 KO, ZNF335 KO, or NT control. Peak validation accuracy (0.70) was achieved at epoch 21. (**C**) Receiver operating characteristic curve of optimized model. Distinguishes true positives (y-axis) vs false positives (x-axis) with a calculated AUC of 0.95. (**D**) Correlation histogram indicating the similarity for each trained cell (x-axis) to one of three conditions (−1 = RNF26 KO, 0 = NT Ctrl, +1 = ZNF335 KO). Cells are colored by their true knockout condition (light orange = RNF26 KO; red = NT Ctrl; purple = ZNF335 KO). (**E**) Confusion matrix depicting the predicted label (x-axis) of each cell in the validation set compared to its true label (y-axis), based on rounding its similarity score to the nearest integer. (**F**) Top scoring CRISPR KO genes cluster into three distinct groups. Gene-level centroid embeddings of the top 358 genes (Z-Score > 2) were extracted before UMAP dimensionality reduction. K-means clustering partitions genes into three phenotypic clusters: cluster 1 (ZNF335-like), cluster 2 (RNF26-like), and cluster 3 (NT-like). (**G**) Hierarchical clustering heatmap showing phenotypic similarity of the top 50 gene knockouts to RNF26 or ZNF335 KO anchors. Darker colors indicate greater similarity to the corresponding anchor (RNF26 KO, left; ZNF335 KO, middle; NT control, right). The RNF26 and ZNF335 KO conditions are labeled in green, while the non-SMC condensin II subunits are labeled in orange.

After optimization, the model correctly classified 70% of individual cells to their respective anchor conditions (Figure 4B), demonstrating strong performance metrics (area under the curve [AUC] = 0.95, loss = 0.14, true vs. predicted correlation = 0.76; Figures 4C, S5B–C). Notably, fewer than 2% of RNF26 KO cells were misclassified as ZNF335 KO and vice versa (Figure 4E), demonstrating the model’s ability to discriminate and report biologically meaningful results. To confirm that predictions were driven by biologically interpretable morphological features, we employed post-hoc saliency mapping based on the integrated gradient (see methods, Figure S5F, S5G).^69^ Consistent with this, the RNF26 KO cells resulted in a higher number of smaller, more heterogeneous nuclear condensates when compared to the ZNF335 KO condition (Figure S5G).

Independent of the model’s continuous regression output, we extracted raw feature embeddings for each CRISPR hit with a Z-score > 2 and averaged single-cell embeddings per condition to generate centroid vectors. These were projected into two dimensions via Uniform Manifold Approximation and Projection (UMAP), ^70^ revealing three distinct clusters by KMeans clustering (Figures S5D, S5E). The resulting UMAP visualization (Figure 4F) separated genes into groups phenotypes resembling the ZNF335 KO cells (cluster 1), RNF26 KO cells (cluster 2), or the NT control cells (cluster 3).

Finally, centroid feature vectors were used to compute the similarity metrics for each gene against the RNF26 KO, ZNF335 KO, and NT control conditions. We clustered gene-level similarity profiles using hierarchical clustering and constructed a heatmap of the top 50 genes sharing high phenotypic similarity to RNF26 KO or ZNF335 KO conditions (Figure 4G). Of interest, knocking out any of the non-SMC subunits of the condensin II complex (NCAPG2, NCAPD3, and NCAPH2)^71^, and to a lesser extent, the condensin II SMC2 subunit, displayed condensates with high phenotypic similarity to the ZNF335 KO condition (Figure 4G).

### RNF26 and ZNF335 Knockouts Exhibit Distinct Phenotypes but Share Accumulation of K48-Ubiquitin

To validate the deep learning results suggesting that condensates arising from knocking out RNF26 and ZNF335 are phenotypically distinct, we used immunofluorescence staining to determine if these condensates accumulate polyubiquitinylated proteins. Similar to those observed in torsin-deficient cells (see chemical screen), condensates in both RNF26 KO and ZNF335 KO cells displayed co-localization with K48-Ub using immunofluorescence imaging (Figure 5A). Next, we assessed whether both condensate conditions co-localize with nucleoporins by performing immunofluorescence staining using α-mAb414, which detects NPC-localized nucleoporins. RNF26 KO cells showed prominent MLF2-GFP co-localization with mAb414 (Figure 5B), analogous to that observed in torsin-deficient cells with stalled NPCs (Figure 5B). In contrast, analysis of confocal Z-stacks revealed that condensates in ZNF335 KO cells rarely shared significant co-localization with mAb414 staining (Figure 5B, 5C), supporting the prediction that RNF26 and ZNF335 KO conditions promote condensates with distinct properties.

**Fig. 5.**
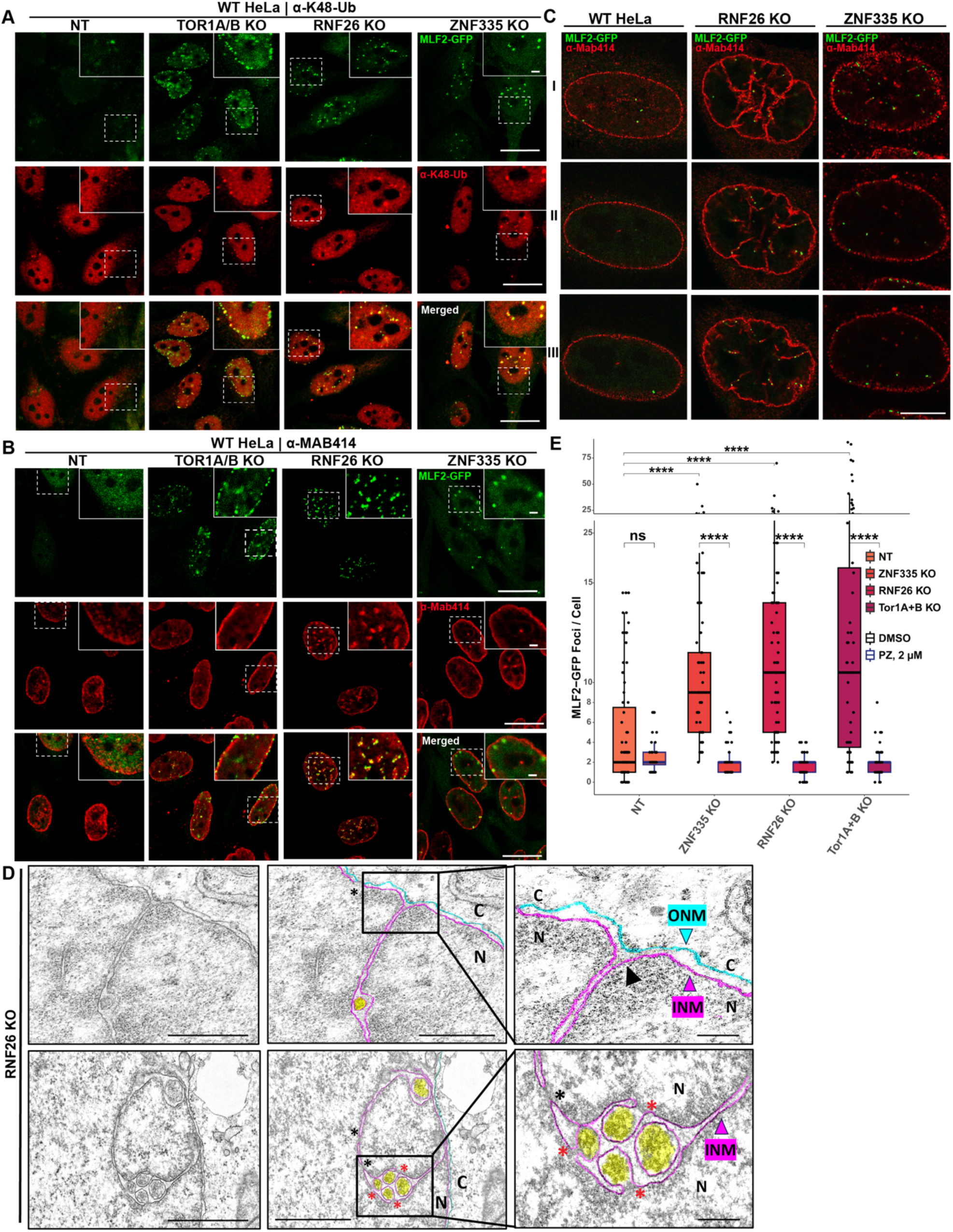
Knockout of RNF26 or ZNF335 induces nuclear condensate accumulation. (**A**) K48-ubiquitinylated proteins accumulate within nuclear condensates. Immunofluorescence confocal images comparing co-localization of MLF2-GFP (green) with α-K48-Ub staining (red) in Cas9-integrated HeLa cells transfected with either (from left to right) NT, TOR1A/B, RNF26 or ZNF335 sgRNAs. (**B**) Condensates of RNF26 KO cells co-localize with NPC markers. Both RNF26 and TOR1A/B KO conditions demonstrated strong co-localization (yellow) of MLF2-GFP (green) with α-Mab414 staining (red). (**C**) Z-stack confocal projections of cells treated with the same conditions as (B). (**D**) TEM images of RNF26 KO cells depicting pronounced nuclear invaginations. Condensates are shaded yellow, while the INM and ONM are outlined in magenta and cyan, respectively. Black arrows draw attention to the interconnectedness of finger-like nuclear protrusions with INM; black asterisks indicate putative NPCs; red asterisks indicate omega-shaped herniations N, nucleus; C, cytoplasm. (**E**) Quantification of MLF2-GFP foci in CRISPR KO cells after PZ treatment. HeLa expressing MLF2-GFP (green) were treated with NT, Tor1A/B, RNF26, or ZNF335 sgRNAs along with either 2 µM PZ or DMSO for 3 h. KO conditions treated with DMSO are shown on the left (black outline) while those treated with PZ are shown to the right (blue outline). Asterisks indicate Bonferroni-corrected p-values by Mann-Whitney U testing (****: p <= 0.0001). IF scale bars at 10 µm (main) and 2 µm (inset); EM scale bars at 250 nm (main) and 100 nm (inset).

Consistent with immunofluorescence data, TEM imaging demonstrated that RNF26 KO cells contain condensates between the nuclear envelope (Figure 5D, shaded yellow). Additionally, RNF26 KO cells feature pronounced NE invaginations reminiscent of nucleoplasmic reticulum formation (Figure 5D).^72,73^ While we observed several NE invaginations contiguous with the inner nuclear membrane (INM), further electron tomographic studies are needed to determine if all invaginations are contiguous with the NE. We then treated RNF26, ZNF335, or TOR1A/B depleted cells with 5% 1,6-hexanediol for five minutes. Only 1,6-hexanediol, but not DMSO or 2,5-hexanediol, dissolved all three types of condensates (Figure S6). Together, these results indicate that while both RNF26 and ZNF335 have a role in safeguarding against nuclear condensate accumulation, only lack of RNF26 results in a condensate phenotype similar to the one observed upon torsin deletion.

### Chemical hit Pyrithione Zinc Modulates Condensates Induced by Genetic Knockouts

Finally, we tested whether compounds identified in our chemical screen could modulate the condensate phenotypes arising from the genetic perturbations, including those linked to microcephaly. Due to the ability of PZ to potently modulate MLF2-GFP condensate markers in torsin-deficient cells (Figures 2D, 2E), we tested whether it could similarly modulate NE condensates in RNF26 KO and ZNF335 KO cells. In both KO conditions, treatment with 2 µM PZ significantly reduced accumulation of nuclear MLF2-GFP condensate foci (Figures 5E, S7). Thus, FDA-approved drugs, including PZ, can be repurposed to modulate condensates across phenotypically distinct genetic perturbations.

## Discussion

This study establishes a scalable high-content platform optimized for identifying chemical and genetic modulators of aberrant condensates that are increasingly implicated in neurological disorders.^12,14,74^ We paired a chemical screen in a DYT1 dystonia disease model with a genome-wide CRISPR KO screen in WT cells enabling complementary analysis of condensate modulation. This integrated approach identified FDA-approved compounds that alter the properties of aberrant condensates in a disease model and uncovered genes that are tied to nuclear condensate homeostasis in WT cells. Notably, several genes identified through this study are mutated in patients with microcephaly, dystonia, and related neurodevelopmental disorders, highlighting previously unappreciated commonalities in the cellular pathology underlying these conditions. In addition, our study identifies previously unknown condensate-modifying genes which may inform both patient-based sequencing and mechanistic studies.

Using the known proclivity of MLF2 to immerse into aberrant condensates,^1^ we exploit MLF2 derivatives as phenotypic condensate reporters. Remarkably, our reporter allows for detection of DYT1 dystonia-associated condensates under single torsin deletion conditions (Fig 3B) that remain undetectable by conventional markers.^28^ In addition, we generalize the utility of MLF2-based reporters for stress granules linked to ALS/FTD (Figure 1B). Notably, MLF2 was recently found as resident in Poly-GA and phosphorylated-TDP-43 inclusions in C9orf72 ALS/FTD patient samples,^75^ as well as co-localization of its *Drosophila* homolog (dMLF) with polyglutamine aggregates^76,77^, suggesting that our readout can be applied to a variety of neuropathological conditions in the future.

Importantly, our screening approach is target-agnostic, utilizing MLF2 solely as a readout for condensate detection at low expression levels to avoid altering condensate dynamics. This allows our platform to be readily adapted to support a range of condensate screening campaigns, even when the affected disease allele is not itself a condensate constituent. This is exemplified by TOR1A/B or ZNF335 KO conditions, which induce condensates via indirect mechanisms, representing a key advantage since condensates are often studied under conditions of disease-allele overexpression, which may perturb physiological condensate equilibria.

Inspired by recent condensate-modifying (c-mod) campaigns,^34,35,78,79^ we screened drugs that could modulate condensates either directly or indirectly. Several of our identified drug hits caused robust re-localization of both our MLF2-GFP biomarker and endogenous (K48-Ub) nuclear condensate markers (Figures 2E, 2F), without significantly altering reporter expression or non-specifically altering physiological phase-environments (e.g., PML bodies, Figure S2G, S2H). To assess the ability to repurpose our chemical hits for other condensatopathies, we compared our top 20 drug hits with two recent screens that identified modulators of stress granules.^34,35^ Half of our drug hits (10 out of 20), including PZ, celastrol, and disulfiram were identified among the top 100 hits of these stress granule screens (Table S4).^34,35^ The ability of identical chemical matter to modulate disparate disease phenotypes underscores the potential of shared etiological principles across aberrant condensates.

To address the incomplete understanding of the dynamics driving condensate formation and the paucity of reliable targets, we repurposed our readout to conduct a genome-wide CRISPR KO screen toward target-directed approaches. Using a deep residual convolutional neural network, further analysis of the resulting condensate phenotypes identified two distinct clusters around either the RNF26 KO or ZNF335 KO condition (Figure 4F). To distinguish these clusters, we performed downstream confocal and electron microscopy on the RNF26 KO and ZNF335 KO conditions. Previous work has implicated RNF26 in ER positioning, endosome retention, and Sec62-mediated recovERphagy.^55,56,80^ While condensates in both knockout conditions accumulated K48-linked ubiquitin (Figure 5A), indicating dysregulation of cellular protein quality control, only RNF26-deficient cells showed strong condensate colocalization with nucleoporins (Figure 5B, 5C) and prominent omega-shaped inner nuclear membrane invaginations (Figure 5C, 5D).

The identification of RNF26—previously unrelated to Torsins or NE morphology—strongly validates our unbiased genetic approach. Building on our prior discovery that MLF2 is a resident of nuclear envelope blebs,^20^ the hallmark phenotype of DYT1 dystonia,^1,19,20,28,29^ we found here that RNF26 deletion provokes a nearly identical cellular pathology characterized by accumulation of MLF2, nucleoporin-associated nuclear condensates, and omega-shaped inner nuclear membrane invaginations, all of which have previously been observed across dystonia models.^1,19,20,28,29,62^ Thus, our study strongly motivates future efforts to explore how RNF26, Torsins, and NE dynamics are functionally connected.

Depletion of ZNF335, a C2H2-type zinc finger and component of the trithorax H3K4-methylation complex,^48^ resulted in the most pronounced condensate phenotype, with the average number of condensate foci exceeding that found in either the RNF26 or TOR1A KO conditions (Figure S4D). Mutations or aberrant splice variants of ZNF335 result in severe microcephaly and frequently reduced life expectancy.^48–53^ Moreover, ZNF335 has been reported to regulate the REST/NRSF complex with its knockdown inhibiting neuronal differentiation. ^48^

In addition to ZNF335, pathogenic allelic variants of 15 out of the top 50 gene hits are associated with neurodevelopmental disorders according to human test results from OMIM^45^ and the NIH GTR.^46^ (Figure 6A). Strikingly, 7/15 of these genes (ZNF335 [MCPH10], CIT [MCPH17], NCAPD3 [MCPH22], CDK5, SIN3A, SHH, and SPDL1) are linked to microcephaly, holoprosencephaly, or lissencephaly (OMIM, GTR), along with additional findings implicating NCAPG2, NCAPH2, and EP300 with microcephaly.^48,64,81–84^ Given that whole-exome sequencing fails to identify causal disease-causing mutations in more than 70% of microcephaly patients,^85^ our curated list of condensate-associated genes provides a valuable resource to prioritize candidates in future patient genetic studies.

**Fig. 6.**
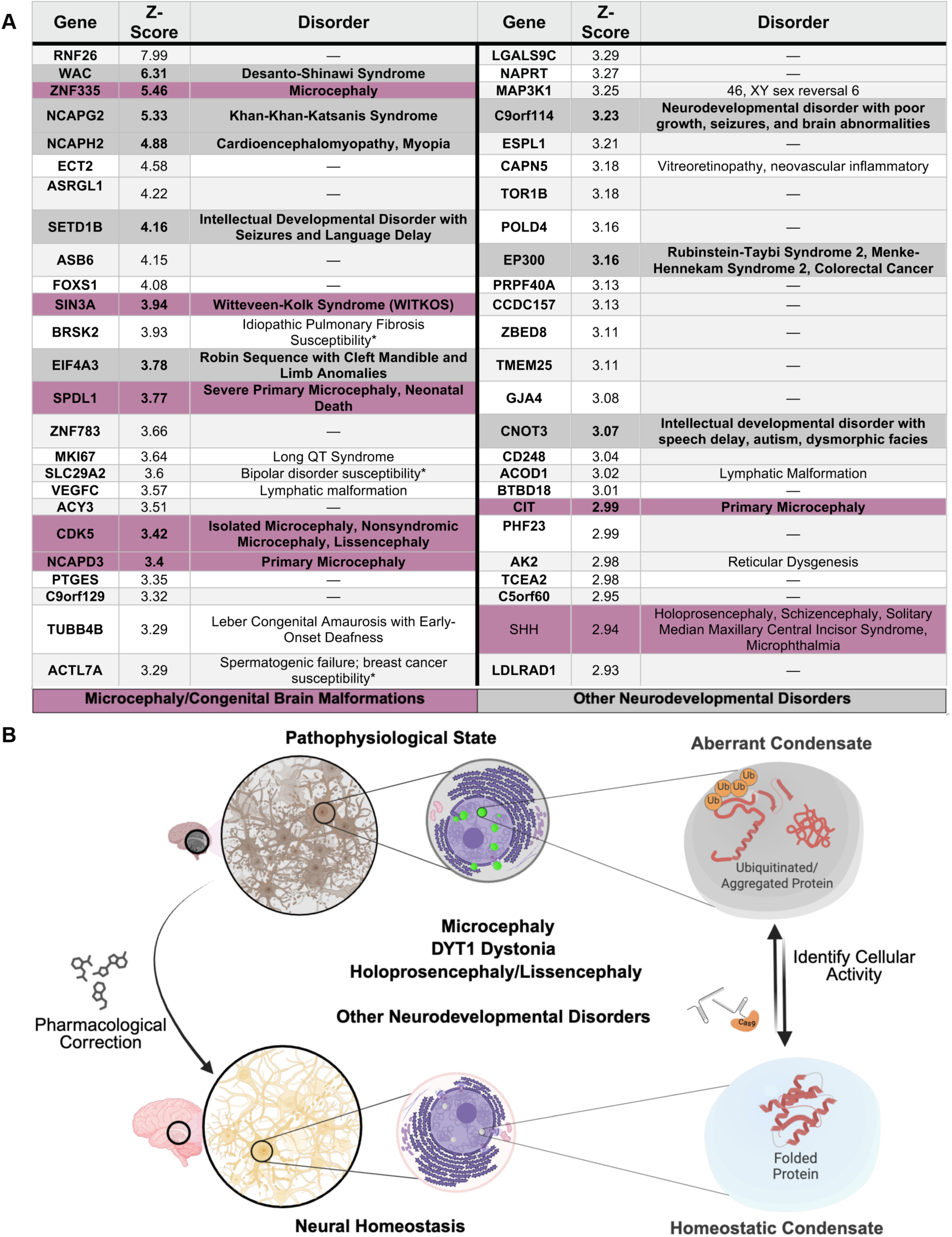
Targeting aberrant condensates as a therapeutic approach for neurodevelopmental disorders. (**A**) CRISPR KO hits enriched for genes implicated in neurodevelopmental disorders. Top 50 candidate genes (left columns), their significance score (Z-Score, middle columns), and associated conditions (right columns) are color-coded based on disorders. Rows labeled purple indicate established link to microcephaly/congenital brain malformations with dark gray indicating additional neurodevelopmental disorders. Genes denoted with an asterisk indicate evidence from genome-wide association studies. Gene relevance to disease was determined based on allelic variants obtained from OMIM and GTR. (**B**) Chemical and genetic modulation of aberrant condensates in neurodevelopmental disorders. Genes involved in the etiology of primary microcephaly and encephalopathies were highly enriched in our screen. Condensates can be modulated using small molecules or genetic perturbations. Targeting condensates could potentially lead to preserving neuronal homeostasis. Arrows indicate possible intervention points and potential outcomes.

Several of the microcephaly-associated gene hits identified in our screen are also involved in chromosomal condensation, including several independently identified subunits of the condensin II complex. Microcephaly has previously been labeled as a “condensinopathy”,^64^ not to be confused with condensatopathies, reflecting prior work that established a relationship between chromosomal condensation defects and reduced cerebral cortex size.^64^ Our deep learning model uncovered an unusually similar nuclear MLF2-containing condensate phenotype between cells deficient in ZNF335 and four out of five of the condensin II subunits (NCAPG2, NCAPH2, NCAPD3, and SMC2; Figures 3B, 4G), inspiring future functional studies at the intersection of MLF2, condensin biology, and phase separation.

In conclusion, our study lays the foundation for exploring what cellular role biomolecular condensates have in microcephaly or related neurodevelopmental disorders (Figure 6B). We overcome the scarcity of generic phase-separation markers by establishing MLF2 as a versatile condensate biomarker and report several chemical modulators of aberrant condensates. This reinforces similar chemical-modulator screening as a general strategy to identify therapeutic interventions across condensatopathies. Our screen and analysis pipeline are readily adaptable for screening campaigns targeting diverse condensate phenotypes, including stress granules and other disease-relevant condensates. This study uncovers a connection between altered condensate accumulation and neurodevelopmental disorders, inspiring pharmacological strategies to correct aberrant condensates.

### Limitations of the study

While our study highlights several chemical and genetic modulators of disease-linked condensates, opportunities exist for further refinement. Pyrithione zinc markedly resolved the proteotoxic condensate cargo in torsin-deficient cells, while complete resolution of the underlying condensates was not observed at the tested concentration (Figure S2I). Extended incubation times may result in condensate resolution, but increased cytotoxicity limited this approach. Similarly, while our genetic screen uncovered an enrichment of neurodevelopmentally-linked genes, redundant pathways or those strictly requiring a neuronal context are likely to be underrepresented and we filtered out conditions that resulted in severe cytotoxicity. Large-scale screening of optimized compound libraries and testing genetic redundancy through combinatorial perturbations will likely identify additional condensate modulators. Furthermore, assessing the condensate burden associated with these neurodevelopmental conditions would benefit from future studies in neuronal or in vivo systems.

## Supplementary Figures

**Fig. S1.**
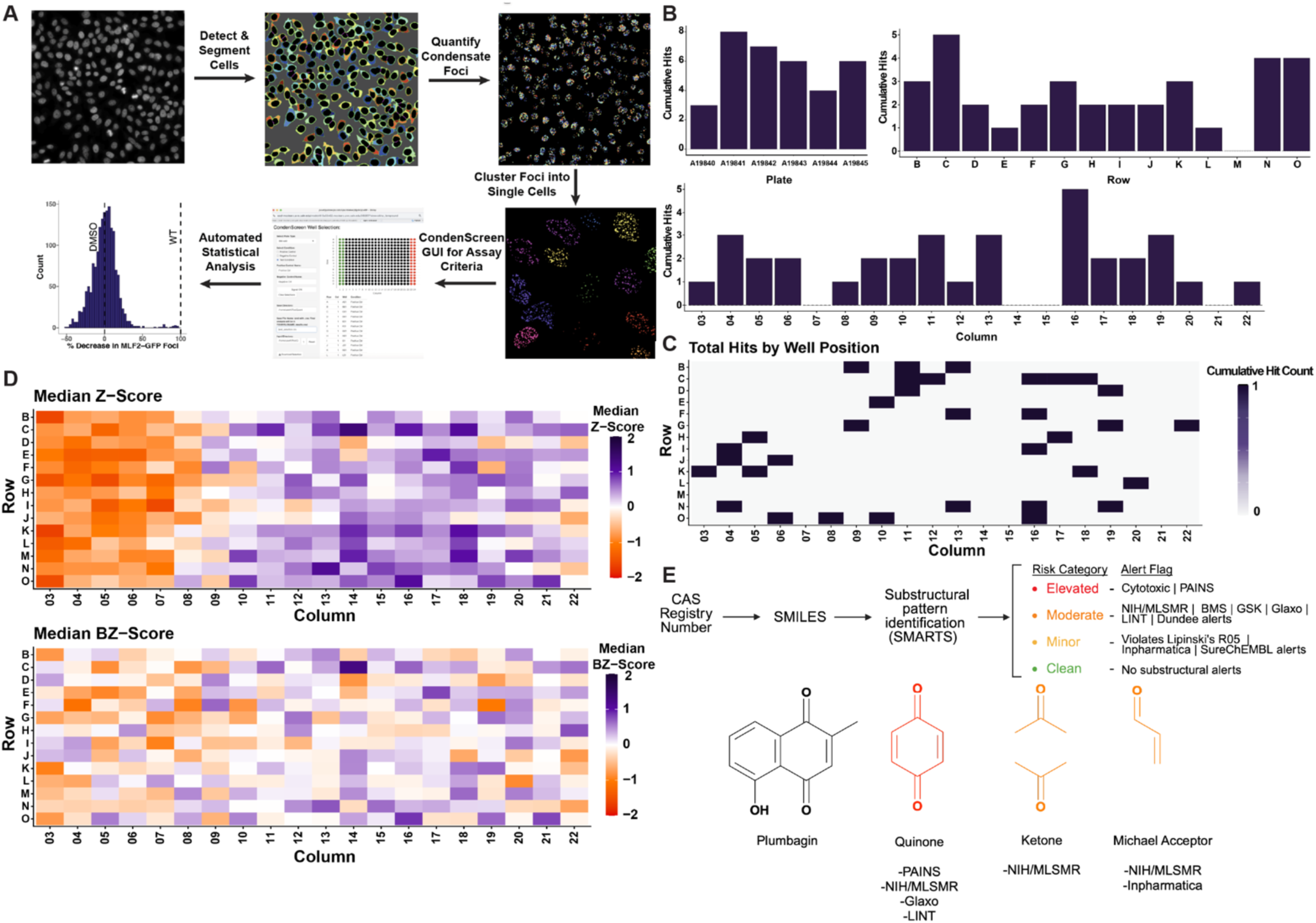
CondenScreen as a high-content condensate analysis pipeline. (**A**) Schematic of the computational *CondenScreen*. Initial cell segmentation and foci detection uses custom *CellProfiler* script and while downstream statistical analysis is completed in *R*. (**B**) Distribution of cumulative hit counts from the Pharmakon 1760 library screen (n = 6 plates) summarized by plate (top left), row (top right), and column (bottom). (**C**) Spatial heatmap of total hits per well position across all plates. Color intensity reflects hit frequency, with darker purple indicating higher cumulative hit counts (a maximum of one hit per well was observed). (**D**) Comparison of hit detection using the Z-Score vs BZ-Score Normalization methods. A heatmap of the median (n = 6) of the Z-Score (top) vs the BZ-Score (bottom) per well. Color indicates score magnitude and direction (orange = negative, purple = positive, white ≈ 0; darker shading reflects larger absolute values). (**E**) Schematic of CondenScreen’s built-in chemical structure hit triage workflow. (top) Using either the name or CAS numbers, the SMILES will be returned (PubChem/NCI CACTUS), followed by substructural pattern identification (SMARTS) flags using several industry alert flags, along with the tested cytotoxicity of each compound. (bottom) Example results using plumbagin. Three alerts were flagged for plumbagin, including a quinone, ketone, and Michael acceptor substructural elements. As quinones are listed as a PAINS substructure, plumbagin received a risk category ranking of “Elevated”.

**Fig. S2.**
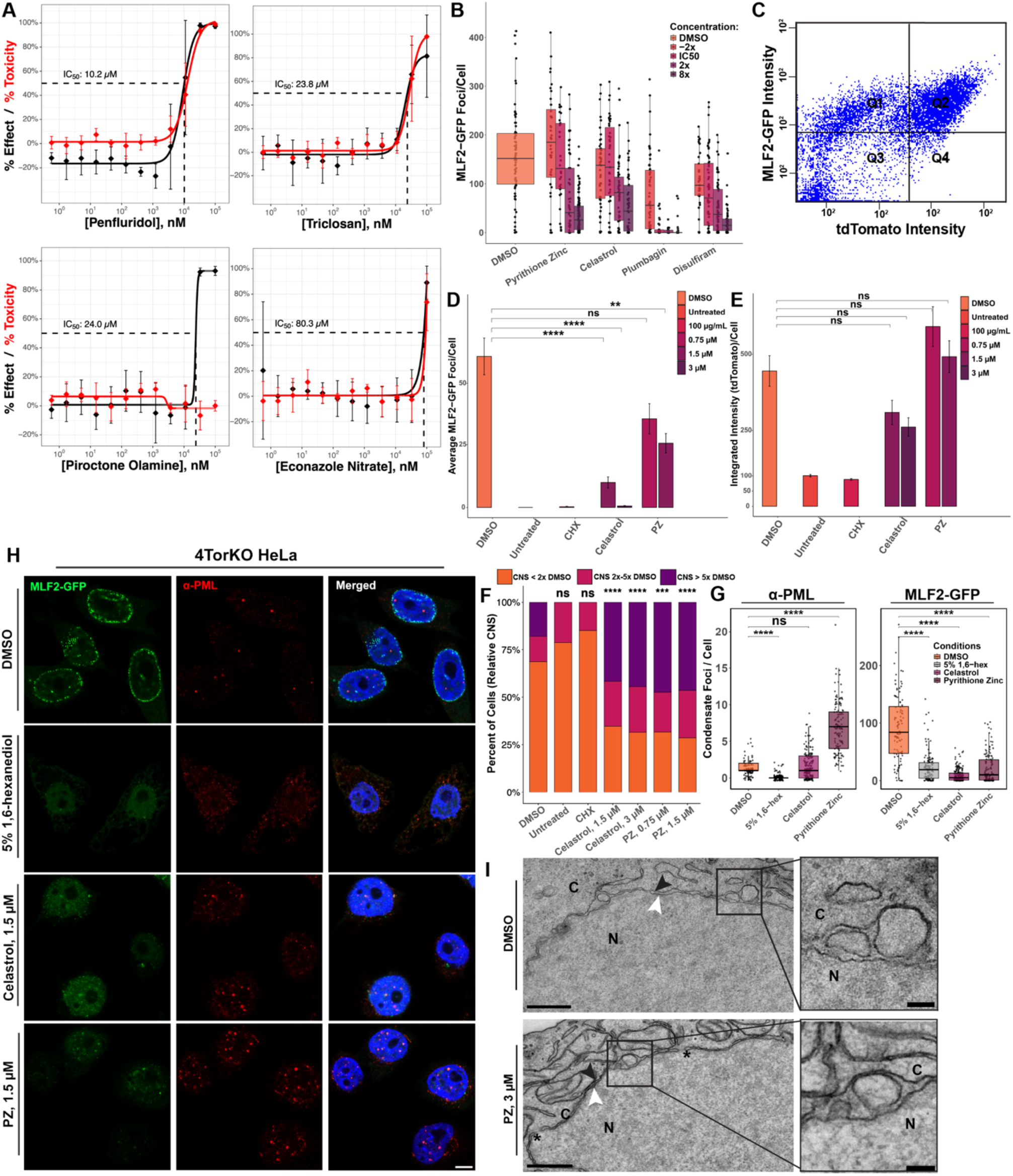
Validation of chemical hits from the Pharmakon 1760 screen. (**A**) Calculated IC_50_ values of additional compounds with dose-dependent relationship. The black and red curves represent % effect and % toxicity, respectively, relative to the DMSO control. (**B**) Drug treatment decreases nuclear MLF2-GFP accumulation. Torsin-deficient cells expressing MLF2-GFP (8 h) were treated at four different concentrations (2x below IC_50_, at IC_50_, 2x above IC_50_, and 8x above IC_50_) for 3 h with either PZ, celastrol, plumbagin, or disulfiram, compared against the DMSO control. Each dot represents the number of MLF2-GFP foci for a single cell, while the horizontal lines indicate the median foci/condition. (**C**) Distribution of FACS results for 4TorKO MLF2-GFP-P2A-tdTomato HeLa cell sorting. Cells within quadrant 2 were sorted. (**D**) Quantification of the number of MLF2-GFP foci/cell (**E**) and tdTomato integrated intensity/cell. Cells (4TorKO) were all (except the untreated condition) induced with Dox and treated with either DMSO, cycloheximide (CHX), celastrol, or PZ. (**F**). Binned CNS value per cell across treatment conditions. Orange indicates <2x the median DMSO CNS value, red indicates 2–5x and purple indicates >5x the median DMSO control. Quantification (**G**) and confocal imaging (**H**) show that PZ does not non-specifically dissolve physiological condensates. Celastrol and PZ reduced MLF2-GFP foci (green) without dissolving PML bodies (red), whereas 5% 1,6-hexanediol dissolved both. All three compounds decreased MLF2-GFP foci relative to DMSO; only 1,6-hexanediol reduced PML foci. PZ increased PML foci number, while celastrol had no significant effect. Scale bars, 5 µm. (**I**) TEM images of 4TorKO HeLa cells treated with either 0.1% DMSO or 2 µM PZ for 3 h. Insets show NE condensates between the inner and outer nuclear envelopes. Scale bars at 500 nm; inset scale bars at 100 nm. Asterisk, putative nuclear pores; black arrowhead, outer nuclear envelope; white arrow, inner nuclear envelope; C, cytoplasm; N, nucleus. Asterisks indicate Bonferroni-corrected p-values determined by Mann-Whitney U testing (**: p <= 0.01; ****: p <= 0.0001).

**Fig. S3.**
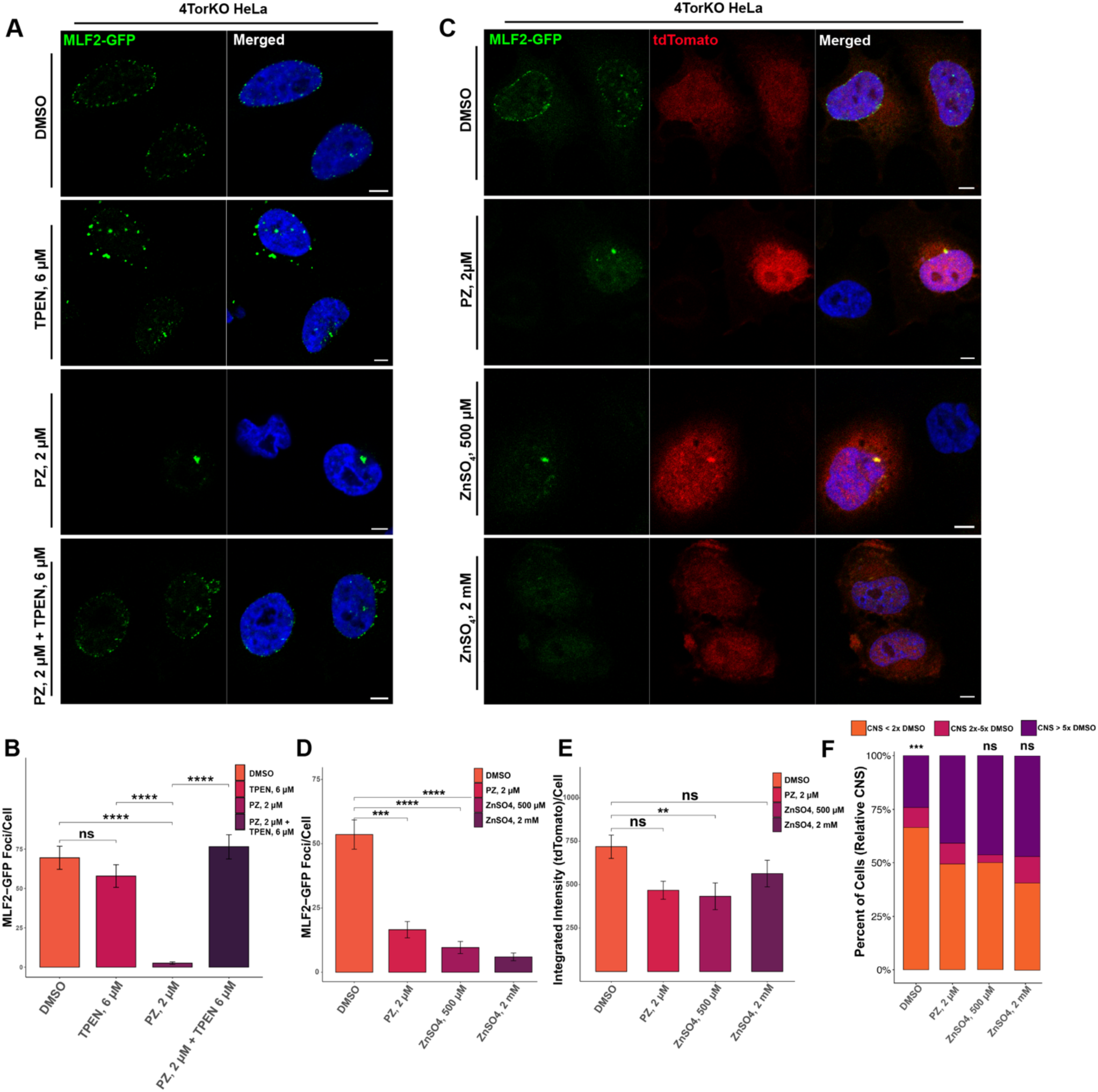
Zinc is sufficient for modulating nuclear condensates in 4TorKO cells. (**A**) TPEN negates the effect of PZ treatment. Confocal images of 4TorKO cells expressing MLF2-GFP (green) and stained with Hoechst (nucleus, blue). Cells were treated with either DMSO, 6 µM TPEN, 2 µM PZ, or 6 µM TPEN + 2 µM PZ (3 h). (**B**) Barplot quantification of (A) showing the average number of MLF2-GFP foci per cell for each condition. (**C**) Zinc sulfate is sufficient to modulate nuclear condensates. Dox-induced 4TorKO were treated with either DMSO, 2 µM PZ, 500 µM ZnSO4, or 2 mM ZnSO4. Representative confocal images demonstrating distribution of MLF2-GFP, tdTomato, and DAPI are shown by the green, red, and blue channels, respectively. The corresponding barplot quantifications of each condition are shown representing (**D**) the average number of MLF2-GFP foci per cell (**E**) the average tdTomato integrated intensity per cell, and (**F**) Binned CNS value per cell across treatment conditions. Orange indicates <2x the median DMSO CNS value, red indicates 2–5x and purple indicates >5x the median DMSO control. Scale bars, 5 µm. Asterisks indicate Bonferroni corrected p-values using a Mann-Whitney U test (*: p <= 0.05; **: p <= 0.01; ***: p <= 0.001; ****: p <= 0.0001).

**Fig. S4.**
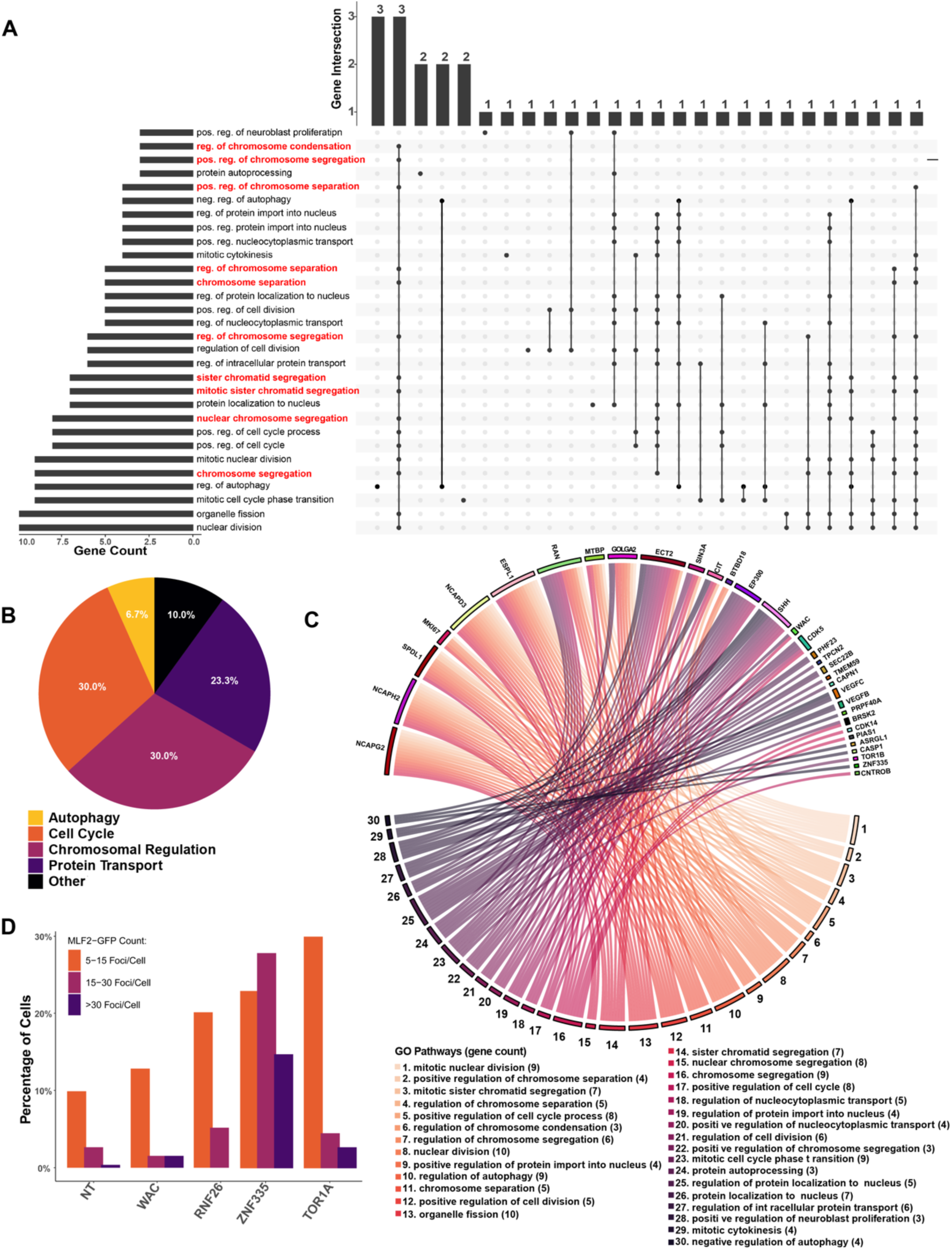
Validation and enriched gene pathways of CRISPR KO hits that promote condensate accumulation. (**A**) Enrichment of pathways related to chromosomal and chromatid regulation. Upset plot depicting relationships between the top 30 GO-enriched biological process pathways. Horizontal bars (left) indicate number of genes for each GO pathway, with terms ranked by gene count. Vertical bars (top) indicate size of each intersecting gene set of genes sharing multiple GO terms. The dot matrix below indicates shared genes among the corresponding GO terms (black dots connected by black line); single dots represent GO terms with unique gene sets. Red dots highlight pathways related to chromosomal or chromatid-related activities, (**B**) Manual clustering of GO biological pathways from the genome-wide KO screen in WT HeLa. The top 30 enriched pathways for the 100 highest-scoring candidate genes were manually clustered within five categories. Percentage represents the fraction of GO pathways within each cluster. (**C**) Chord diagram mapping each gene to its specific GO terms. Each gene (outer labels, top) is linked to its corresponding GO terms (bottom labels), with colored ribbons denoting relationships. (**D**) Binned quantification of the MLF2-GFP foci count after CRISPR KO of select candidate genes. RNF26 KO, ZNF335 KO, and WAC KO HeLa cells were compared against NT control. Bars indicate percent of cells with 5-15 foci MLF2-GFP foci/cell (orange), 15-30 foci (violet), or >30 foci (dark purple).

**Fig. S5.**
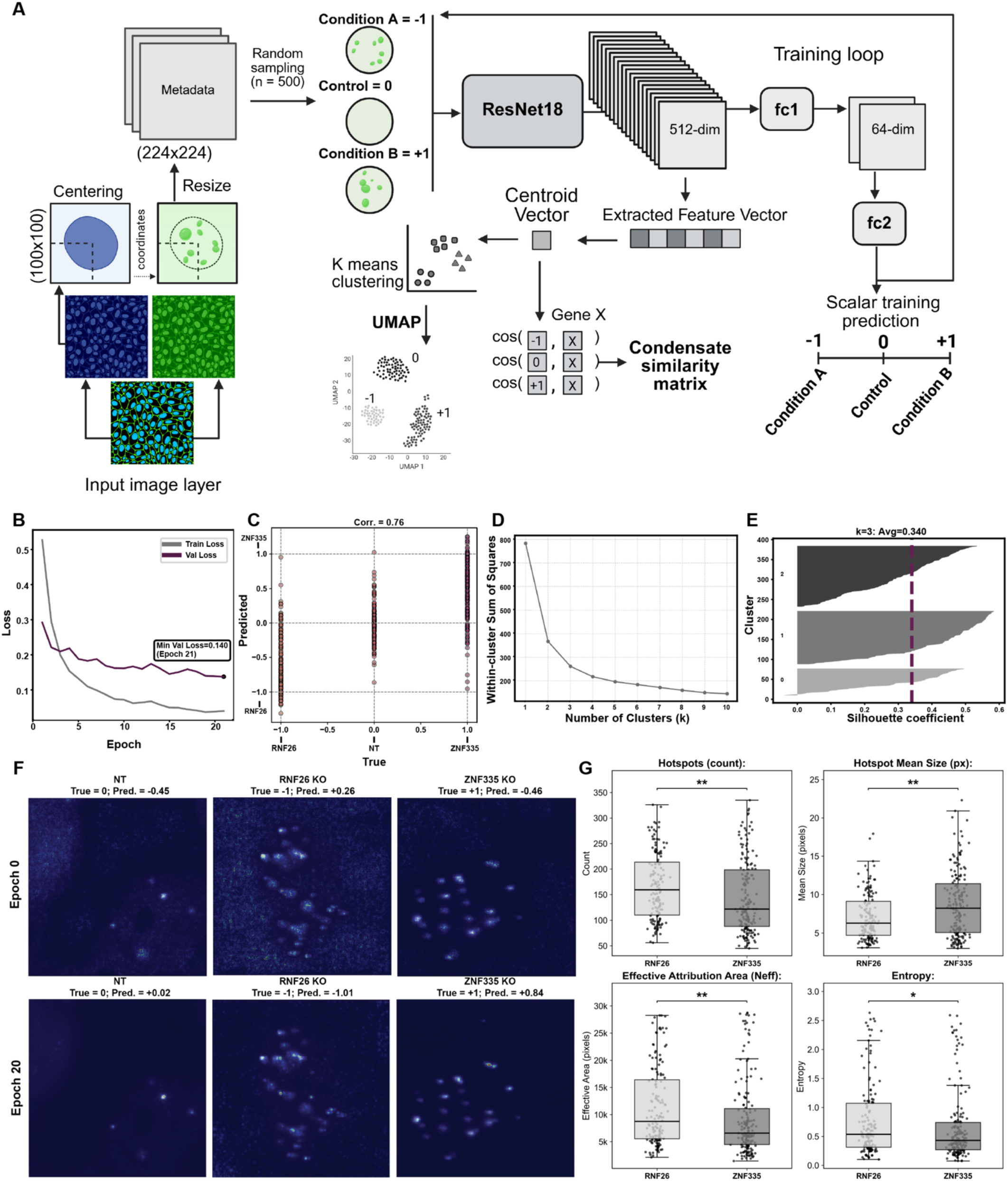
Validation of deep learning model to detect subtle variations in condensate phenotypes. (**A**) Schematic workflow of the deep learning-based condensate phenotype analysis pipeline. The model was trained on raw images from the CRISPR screen, which were resized to single-cell patches and labeled. A residual convolutional neural network (ResNet18) was optimized to distinguish variations in condensate morphology. Centroid feature vectors from the top CRISPR hits (Z-score > 2) were extracted and mapped using UMAP and k-means clustering. (**B**) Training and validation loss curves based on Huber loss (nn.SmoothL1Loss). Loss of each epoch for both the training set (gray) and validation set (purple) are reported with a minimum loss (0.14) occurring at epoch 21. (**C**) Correlation scatter plot of validation set. Each dot represents the similarity score (−1 = RNF26 KO; 0 = NT Ctrl; +1 = ZNF335 KO) of a single cell from the validation set. The predicted value (y-axis) is compared against the true value (x-axis), with a Pearson correlation score of 0.76. (**D**) Elbow (scree) plot showing the within-cluster sum of squares across cluster numbers (k = 1-10). (**E**) Silhouette plot showing the silhouette coefficient (0.34, vertical dashed line) for three clusters. (**F**) Representative saliency maps prior to training (top, Epoch 0) vs post-training (Epoch 20). From left to right, non-targeting control cell (true label = 0, initial pred. = −0.45, final pred. = +0.02), RNF26 KO cell (true label = −1, initial pred. = +0.26, final pred. = −1.01), and ZNF335 KO cell (true label = +1, initial pred. = −0.46, final pred. = +0.84). (**G**) Boxplots of the hotspot count, mean size of hotspots, effective attribution area, and Shannon’s Entropy across the single cells (dots). Asterisks indicate p-values using a Mann-Whitney U test (*: p <= 0.05; **: p <= 0.01).

**Fig. S6.**
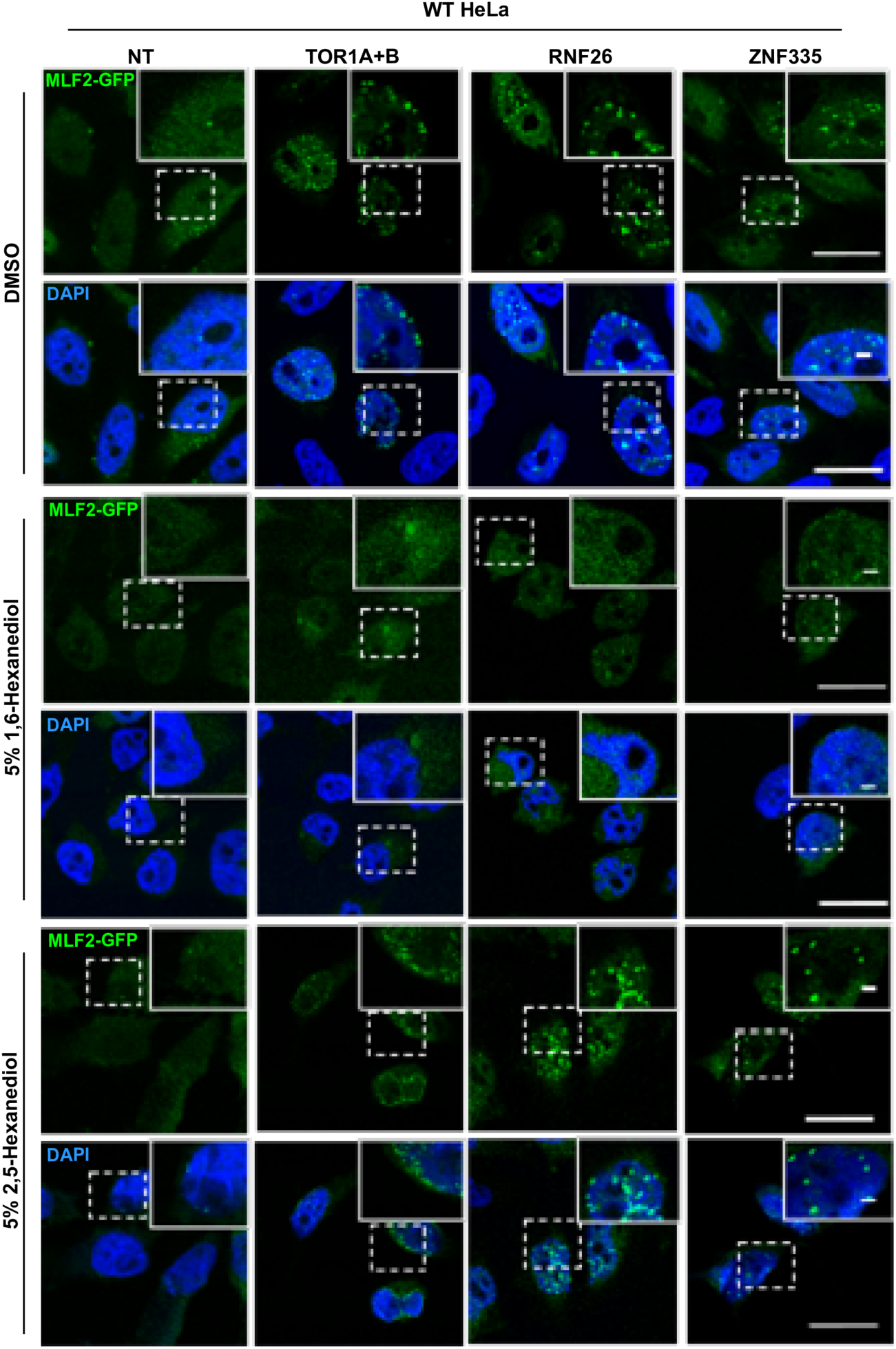
Condensates resulting from CRISPR KO can be dissolved by 1,6-Hexanediol. (A) Fluorescent microscopy images of Cas9-integrated WT HeLa cells treated with sgRNAs against either NT control, TOR1A/B, RNF26, or ZNF335 (from left to right) for 72 h. Cells were then exposed to either DMSO (top), 5% 1,6-hexanediol (middle), or 5% 2,5-hexanediol (bottom) for 5 min. WT cells with ZNF335, RNF26, or TOR1A/TOR1B knocked out demonstrated dissolution of condensates marked by MLF2-GFP (green) only after exposure to 5% 1,6-hexanediol (middle), but not DMSO (top) or 5% 2,5-hexanediol (bottom) controls.

**Fig. S7.**
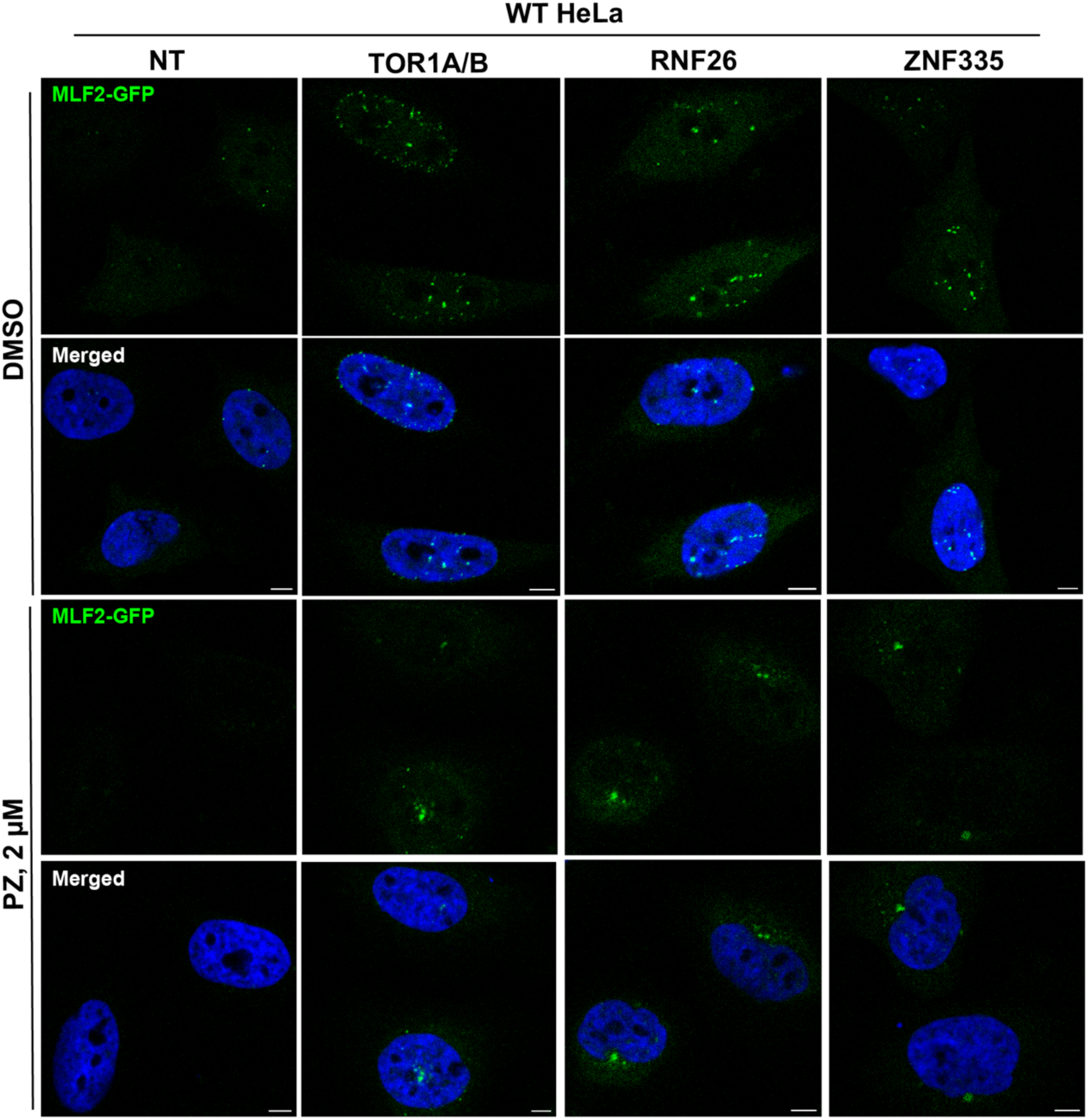
Pyrithione zinc modulates atypical nuclear condensates across genetic perturbations. Cas9-integrated WT HeLa cells treated with sgRNAs against either NT control, TOR1A/B, RNF26, or ZNF335 (from left to right) for 72 h. Cells were then induced with MLF2-GFP (6 h Dox) and treated with either 0.1% DMSO (top panel) or 2 µM PZ (bottom panel) for 3 h. Representative confocal images show MLF2-GFP (green) and DAPI (blue). Scale bars, 5 µm.

## Materials and Methods

### Tissue culture

Adherent cultured cell lines (HeLa and HEK293T) were maintained in Dulbecco’s modified Eagle’s medium (DMEM) supplemented with 10% (vol/vol) heat-inactivated fetal bovine serum (Thermo Fisher Scientific) and 100 units (U) /mL penicillin-streptomycin (Thermo Fisher Scientific). Cells were maintained in either T75 flasks or 10 cm dishes at 37°C (95% humidity; 5% CO_2_) and passaged every third day. For splitting, the culture medium was removed, washed once with 1x PBS (Gibco), trypsinized with 0.25% trypsin-EDTA (Gibco), and transferred to new dishes (diluted 1:5 or 1:10).

### Plasmid Constructs

Unless otherwise stated, all plasmid constructs were subcloned into a pcDNA™3.1 (+) Mammalian Expression Vector (Invitrogen, V79020) using standard Gibson Assembly protocols. The cDNA for the MLF2-GFP-P2A-tdTomato construct (MLF2 GenBank accession no. <u>BC000898</u>) was subcloned into the pRetroX-Tight-Pur-GOI (gene of interest) vector (Takara Bio) and co-transfected with a pRetroX-Tet-On advanced inducible expression construct (Takara Bio). Viral constructs were co-transfected with both a VSV-G viral envelope plasmid (Addgene #12259) and an MMLV gag/pol viral packaging plasmid. For generation of HeLa cells expressing Cas9-mCherry, an mCherry tag was inserted into the lentiviral plasmid containing Cas9-Flag (Addgene #52962) immediately after the Flag tag into the BamHI cloning site.

### Viral Production

Lentivirus production was carried out as previously described.^86,87^ Briefly, to generate Cas9-flag lentivirus, low-passage HEK293T cells were plated in a 10 cm dish with a density of 3.5×10^6^ cells per dish. Medium was replaced the next day with 6 mL fresh (antibiotic-free) DMEM and transfected with three plasmids encoding: psPAX2 (Addgene #12260) (2 μg), viral envelope protein VSV-G (1 μg), and lenti-Cas9-Flag-mCherry (6 μg) using Lipofectamine 2000 (Thermo Fisher Scientific) at a 3:1 ratio of DNA to Lipofectamine. After 72 h, supernatants containing the virus encoding Cas9-mCherry were harvested and filtered through a 0.45 μm Steriflip filter unit (MilliporeSigma), aliquoted into screw-cap tubes, and stored at −80°C or used for transduction immediately. For the Dox-inducible MLF2-GFP-P2A-tdTomato system, two separate retroviruses were produced using three plasmids encoding: MMLV gag/pol (2 μg), viral envelope protein VSV-G (1 μg), and either pRetroX-Tight-MLF2-GFP-P2A-tdTomato or pRetroX-Tet-On (6 μg) using Lipofectamine 3000 (Thermo Fisher Scientific) at a 2:1.5 ratio supplemented with Lipofectamine P3000 (2.5:1). After 72 h, supernatants were collected, stored, or used, as described.

### Generation of stable HeLa cell lines

The detailed protocol regarding the Tor1A/1B/2A/3A KO and integration of a Dox-inducible MLF2-GFP reporter in HeLa cells has been extensively described by our lab (Laudermilch et al. 2016; Rampello et al. 2020).^20,28^ The Cas9-flag-mCherry HeLa and Dox-inducible MLF2-GFP-P2A-tdTomato 4TorKO HeLa cell lines were generated by seeding Dox-inducible MLF2-GFP WT HeLa or 4TorKO cells in six-well plates. The next day, media was replaced with DMEM (antibiotic-free) supplemented with 4 μg/mL polybrene (Sigma-Aldrich), and 100 μl of either the Cas9-Flag-mCherry virus (WT HeLa) or both 100 µL of the MLF2-GFP-P2A-tdTomato and pRetro-Tet-On viruses (4TorKO) added dropwise to wells. Medium was replaced 48 hours after transduction (complete DMEM) and grown for another 24 hours. Cells were collected in sorting buffer (ice-cold 1x PBS, Gibco) and filtered through a 40 μm nylon mesh filter. Using a BD FACS Aria III, cells were sorted as single cells by selecting those high in either mCherry (WT HeLa) or MLF2-GFP/tdTomato (4TorKO, 16 h after 0.5 µg/mL Dox induction) fluorescence intensity. A stringent gating strategy was applied to enrich for cells positive for the respective fluorophores while excluding those with extremely high signal intensity, to obtain a pure yet physiologically representative population.

### High-content phenotypic screening strategy

All high-content screening was performed at Yale’s Center for Molecular Discovery (YCMD). The Horizon Discovery Edit-R synthetic sgRNA library was used in the whole genome CRISPR KO screen, targeting 19,195 human genes, and comprising of the Drug Targets library (3,683 genes, GA-004650-E2, Lot 20126), the Druggable Genome library (4,643 genes, GA-004670-E2, Lot 20120), and the Human Genome library (10,869 genes, GA-006500-E2, Lot 20130). Each screening plate included 16 replicates of the negative control (NT sgRNA) and 16 replicates of the positive control (combined Tor1A and Tor1B sgRNAs). To ensure robust KO, a pool of three sgRNAs targeting the same gene at different regions was pre-plated into each arrayed well of 384-well glass-bottom plates (Cellvis, #384-1.5H-N). Addition of sgRNA to screening plates was done using the Labcyte Echo 550 Acoustic Dispenser (Labcyte, now Beckman) at a final concentration of 25 nM.

Reverse transfection was performed using DharmaFECT1 Transfection Reagent (Horizon/Dharmacon), which was diluted at 1:500 in Opti-MEM (Thermo Fisher Scientific) and mixed with sgRNA. The mixture of sgRNA and transfection reagent was incubated for 30 minutes, after which, 2,000 cells/well (WT HeLa integrated with Cas9/MLF2-GFP) in antibiotic-free DMEM were added using a Thermo MultiDrop Combi (Thermo Fisher Scientific). The plates were then incubated for 72 hours at 37°C. To induce MLF2-GFP reporter expression, 0.5 µg/mL Dox was added to each well 16 h before fixation. Media was removed and cells washed once with PBS and fixed with 4% paraformaldehyde (PFA) for 20 minutes. Cells were washed twice with PBS, permeabilized with 0.1% Triton X-100 for 5 minutes, and washed again with PBS. Cells were stained with Hoechst 33342 dye (1:1000) (Invitrogen, H1399) and HCS CellMask Deep Red Stain (Invitrogen, H32721) (1:5000 of 10 mg/mL, Thermo Fisher Scientific) diluted in PBS for 30 minutes, and washed a final time in PBS. The Molecular Devices ImageXpress 4.0 high-throughput fluorescent microscope (40x magnification) was used to obtain 9-image sets (each consisting of DAPI, MLF2-GFP, and Deep Red CellMask images) per well, resulting in 10,368 TIFF images/plate. Images were uploaded to Yale’s McCleary high-performance computing cluster (Yale Center for Research Computing) for analysis.

For chemical screening, the Pharmakon 1760 drug library (MicroSource Discovery Systems, Inc.) was used, consisting of 1360 FDA-approved drugs and 400 International drugs on six 384-well plates. Each screening plate consisted of 16 replicates of the negative control (4TorKO HeLa cells treated with DMSO [0.1% v/v]) and 16 replicates of the positive control (WT HeLa cells treated with DMSO [0.1% v/v]). Plates were seeded with 4,000 HeLa cells/well (Multidrop Combi) in 20 µL of DMEM before incubating at 37°C for 24 h. MLF2-GFP expression was induced by adding 0.5 µg/mL of Dox into each well 8 h prior to fixation. Compounds were resuspended in DMSO and added to each well 2 h prior to fixation using a Labcyte Echo 550 Acoustic Dispenser (Labcyte, now Beckman), at a final concentration of 10 µM. Cells were washed, fixed, stained, and imaged using the methods outlined above.

### Image segmentation of phenotypic screen data

We created a custom analysis pipeline to segment condensate foci into single cells using the open-source software *CellProfiler 4.2.6.*^38,39^ and downstream statistical analysis using *R Studio* (Version 2023.12.1+402).^88^ Individual nuclei were distinguished by shape for objects between 30-100 (+4) pixels using an adaptive (40 pixels) Minimum Cross-Entropy thresholding (0.01-1.0 threshold) in *CellProfiler 4*. The cytoplasm of each nucleus was delineated as a secondary object of each nucleus using a watershed algorithm with global Otsu thresholding (three classes, foreground; 0-1.0 threshold, threshold correction factor = 0.7).^39,89^ Each GFP-channel image was enhanced using a white top-hat transform (speckles; maximum 10 pixels), allowing each condensate foci between 1-10 pixels to be detected using adaptive (50 pixels) Otsu thresholding (three classes, foreground; 0.0175-1.0 threshold, threshold correction factor = 1.2). Groups of condensate foci were then distinguished using local intensity and related to their corresponding cytoplasm/nuclei. The same image analysis pipeline was applied across all imaging experiments, with only minor adjustments (e.g., cell size or foci intensity thresholds). Any parameter changes were applied uniformly across all conditions and controls within a given experiment.

For parallel analysis, a Bash script was developed allowing the execution of multiple headless CellProfiler instances on a Simple Linux Utility for Resource Management (SLURM)-managed compute cluster. Specifically, after constructing the analysis pipeline in the CellProfiler GUI, the CreateBatchFiles module is used to generate a batch configuration file that defines image metadata and processing instructions. This batch file is then imported to the cluster environment, and all images are registered within a single CellProfiler project in array format, with images from each plate corresponding to a separate SLURM job. This approach enables autonomous analysis of over 1 million high-content images. TIFF images—amounting to multiple terabytes of data—within a matter of hours. Both the CellProfiler pipeline and Bash scripts are open source and available publicly on the SchliekerLab GitHub page as part of the CondenScreen pipeline (https://github.com/SchliekerLab/Condenscreen.git).

### Statistical analysis of screen data

We developed a condensate screening using a combined *CellProfiler* and *R* pipeline, *CondenScreen*, to statistically analyze all screen data. The *R*-script features an interactive GUI allowing the user to select plate size (6-well to 384-well format), plate layout, and input whether the data was taken from a signal “ON” or a signal “OFF” screen. With limited user input, it then automatically iterates through images from each plate and assigns groups of image sets into their corresponding wells using string parsing. To control for variations in transfection efficiencies that could inadvertently impact downstream scoring, the program has an optional filter to select only the top 25% of cells showing the highest (or lowest) signal using the modest assumption that at least 1 out of 4 cells, on average, will have received the sgRNAs of interest. Additionally, for both screens, the image with the highest score was filtered out, as there were several conditions where one image-set had screening aberrations skewing the condition toward a false positive result. For each plate, aggregate statistics of the respective positive and negative controls are compared, including signal-to-background (SB), standard deviation (SD) of the controls, and the coefficient of variation (CV) of the controls, among others. To determine separation between positive and negative controls, a Z’ assay quality score was calculated: *Z*′ = 1 − (3(*σ_PC_* + *σ_NC_*)/|*μ_PC_* − *μ_NC_*|),^40^ where *σ_PC_* and *σ_NC_* is the SD and *μ_PC_* and *μ_NC_* is the condensate mean of the positive and negative controls, respectively.

The percent effect of each condition was quantified by either 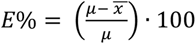 (for signal “OFF” selection) or 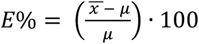 (for signal “ON” selection) where 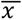 is the average number of condensate foci/cell per tested condition, and *μ* is the average number of condensate foci/cell across the screening population of the plate. The significance scoring function is automatically changed depending on selection of a signal “ON” vs a signal “OFF” screen, although this can be adjusted.

For signal “ON” screens, the significance of each condition was calculated using the Z-Score to normalize for intraplate variations: *Z* = (*E*% − *μ_E_*_%_)/*σ_E_*_%_, where *μ_E_*_%_ and *σ_E_*_%_ represent the population average percent effect and standard deviation, respectively. Since signal “OFF” screens can often have increased variability, the significance of each well was quantified by calculating the BZ-Score. First, B-Scores were computed by applying Tukey’s median polish to each plate to remove row, column, and edge effects. The residuals from the median polish were then scaled by the median absolute deviation (MAD) to produce a robust per-well B-score. Then, the BZ-Score was calculated using the following formula: *BZ* = (*BScore* − *μ_BScore_*)/*σ_BScore_*, where *μ_BScore_* and *σ_BScore_* are the mean and standard deviation of the B-Scores over the entire plate.^90,91^

To determine how chemical or genetic perturbations alter cell viability, we derived a function to extrapolate differential cellular health by morphological characterization of the nucleus, which was used during validation experiments. Specifically, we added several modules to *CellProfiler* to measure the size, shape, intensity, and texture of each nucleus.^38^ This is crucial as several compound concentrations had a non-significant effect in changing the total cell count, but greatly disrupted nuclear morphology based on visual inspection. After testing the effects of multiple parameters, we derived the following empirical formula which best represented the data to estimate nuclear health: *NucHealth* = Δ*_CC_* + ((100 + Δ*_CC_*)/100) · ∑(Δ*_Area_* + Δ*_FF_*)/2, where Δ*_CC_*, Δ*_Area_*, and Δ*_FF_* are the normalized difference in cell count, nuclear area, and nuclear FormFactor (measurement of nuclear irregularity) compared to the screening population (or DMSO for dose-response), respectively. Normalization was calculated according to: Δ*_ν_* = ((*V* − *μ_V_*)/*μ_V_*) · 100, in which *V* is the variable of interest. The equation was derived with the following assumptions and limitations. (1) Changes to the area and FormFactor should be weighted equally, and (2) cell death should be given a higher impact weight over morphological changes.

Two optional *CondenScreen* quality control features allow for flagging potential false-positives or problematic hits arising from screening large libraries. In addition to detecting spatial intra-plate bias, the program flags screen actives arising from global systematic inter-plate spatial biases. These include shifts in continuous significance score metrics (Z-Scores and BZ-Scores) and enrichment of binary hits across rows, columns, individual well positions, and plates with a statistically enriched hit rate. Continuous metric shifts are assessed using a Wilcoxon rank-sum (Mann–Whitney U) test comparing each group to all other wells, while hit enrichment is assessed using Fisher’s exact test. P-values are Benjamini–Hochberg adjusted across groups and flagged at an adjusted p-value < 0.05.

*CondenScreen* enables flagging potentially problematic chemical structures using compound SMILES (Simplified Molecular Input Line Entry System), either provided manually (recommended) or can attempt to be retrieved automatically from PubChem^92^ using the CAS number, with CACTUS^93^ (NCI Chemical Identifier Resolver, https://cactus.nci.nih.gov/chemical/structure) used as a fallback. Smiles are processed using RDKit (via Reticulate in R), and compounds are evaluated against SMARTS (SMILES Arbitrary Target Specification) substructures appearing in alert libraries compiled from RDKit and ChEMBL resources. We utilized a pre-compiled list of these structural alert filters from a csv document allowing SMARTS identification within the RDKit environment, compiled by Pat Walters (see GitHub for more details: https://github.com/PatWalters/rd_filters.git). We defined compounds as having an “Elevated” risk if they include SMARTS overlapping with the Pan-Assay Interference Compounds (PAINS) alerts,^94,95^ NIH Molecular Libraries Small Molecule Repository (MLSMR) alerts, or have a significant cytotoxicity (≥25% viability loss); “Moderate” risk if overlapping with the Bristol-Myers Squibb (BMS), GlaxoSmithKline (GSK/Glaxo), Pfizer LINT alerts, or the University of Dundee filter sets, or tdTomato reporter suppression (≥25% decrease); and “Minor” risk if violating the Lipinski rule-of-five^96^ or appearing in the Inpharmatica or SureChEMBL medicinal chemistry alert libraries. Compounds lacking these features were classified as “Clean.” These categories are intended to support chemical triage rather than exclude compounds.

*CondenScreen* provides additional downstream inter-plate and intra-plate visualizations using open-source libraries, including *ggplot2*^97^ and *ggplate* (10.32614/CRAN.package.ggplate). The output includes a ranked list of hits, plots of the screening distributions (raw condensate foci/cell and %Effect), a Z/BZ-score plot, and individual plate heatmaps for each 384-well plate. Finally, top hits in both screens were inspected manually to ensure quality control and rule out any additional potential screening aberrations. Notably, manual inspection was rarely a reliable measure of hit robustness. In several cases, hits that were initially overlooked during first-pass screening (false negatives) were later validated and correctly identified by the program. The *CondenScreen* platform is publicly available on the SchliekerLab GitHub page: https://github.com/SchliekerLab/Condenscreen.git

### Development of phenotypic machine learning model to distinguish condensate phenotypes

We developed a supervised machine learning model to differentiate between subtle phenotypic differences in nuclear condensates. The model was built around the ResNet18 architecture^67^ with pretrained initial starting weights borrowed from ImageNet1k IMAGENET1K_V1^68^ using torchvision^98^ on Python 3.11.8 (conda-forge). Using images directly from the CRISPR KO screen, 4,523 single cells were extracted, cropped (100×100) and resized (224×224) before centering (DAPI-channel) each cell. Single cells were labeled in accordance with their file names, with NT Ctrl cells = 0, RNF26 KO cells = −1, and ZNF335 KO cells = +1. To ensure even sampling numbers across conditions, cells from the overrepresented NT Ctrl conditioned were downsampled to match KO conditions, with only the MLF2-GFP channel selected for analysis. Data were split 80:20 into train and validation sets (sklearn.model_selection: train_test_split)^99^. Tensors were packaged (torch.utils.data) before employing the following parameters during model training (PyTorch^100^): AdamW (lr = 1e-4, weight decay = 5e-2), dropout = 0.5, hidden dimension = 64, SmoothL1Loss, CosineAnnealingLR, 100 max epochs, and early stopping (patience = 25). Model performance was evaluated using several modules from sklearn.metrics and scipy.stats,^99,101^ including: mean_absolute_error(); r2_score(); pearsonr(); accuracy_score(); confusion_matrix(); roc_curve(); roc_auc_score(); and ConfusionMatrixDisplay(). The integrated gradient^69^ for the validation data was used for post-hoc saliency mapping in order to ensure the model was learning biological features compared to technical noise. Briefly, for each cell, an attribution map was calculated using integrated gradients, which averages model gradients across 24 interpolation steps between the input image and a zero baseline. This attribution map was min-maxed normalized and converted to a single-cell 2D heatmap. For each cell, the spatial distribution of saliency attribution (Shannon entropy;^102^ heat_entropy), effective number of pixels contributing to the saliency distribution (heat_effective_npix), and hotspot number and mean size (heat_hotspot_components), where hotspots were defined as connected components within the top 2% of saliency intensities. The per-cell distributions for ZNF335 vs RNF26 cells were statistically assessed using a two-sided Wilcoxon rank-sum (Mann–Whitney U) test.

After optimization, all CRISPR KO gene hits with a Z-Score > 2 were cropped, resized, and centered before randomly downsampled to 500 cells per condition (accounting for differences in cell viability). Cells were then analyzed by the optimized model and their latent representation embeddings (512-dimensional latent embeddings per cell, e.g., condensate morphology, size, intensity, etc.) were extracted. Relative centroid vector scores were computed (mean embedding across cells), and KMeans clustering with *k* = 3 was applied to UMAP (n_neighbors = 15, min_dist = 0.1) coordinates to assign genes to discrete clusters. The optimal cluster number was selected using the elbow method (KMeans, k = 1–10) and silhouette analysis (sklearn.metrics: silhouette_samples | silhouette_score), yielding three distinct clusters. The difference in the cosine similarity for each gene’s centroid vector embedding was compared with the three anchor centroids (RNF26/NT Ctrl/ZNF335) and exported to an .xlsx file (see Table S3). A hierarchical heatmap was constructed using the top 50 genes with highest similarity to the RNF26 and ZNF335 KO conditions, via pandas, seaborn.clustermap, and matplotlib.pyplot. For a full list of modules imported and model training parameters, please refer to our source script: https://github.com/SchliekerLab/Condenscreen.git

### Solubilization of hit compounds

All putative hit compounds were resuspended up to 100 mM in DMSO. The following compounds were solubilized: pyrithione zinc (PZ) (CAS 13463-41-7), celastrol (CAS 34157-83-0), nerol (CAS 106-25-2), spiramycin (CAS 8025-81-8), choline chloride (CAS 67-48-1), disulfiram (CAS 97-77-8), bronopol (CAS 52-51-7), penfluridol (CAS 26864-56-2), plumbagin (CAS 481-42-5), dyphylline (CAS 479-18-5), econazole nitrate (CAS 24169-02-6), ethacrynic acid (CAS 58-54-8), piroctone olamine (CAS 68890-66-4), dihydrocelastryl (CAS 80-97-7), fenipentol (CAS 583-03-9), triclosan (CAS 3380-34-5), eprodisate disodium (CAS 36589-58-9), L-phenylalanine (CAS 63-91-2), nabumetone (CAS 42924-53-8), edrophonium chloride (CAS 116-38-1), josamycin (CAS 16846-24-5), mevastatin (CAS 73573-88-3), clofibric acid (CAS 882-09-7), docosahexaenoic acid (CAS 6217-54-5).

### Dose-dependent validation of small molecules

Based on the change in MLF2-GFP foci, viability, and structural diversity, the 22 top-scoring putative hit compounds were resuspended to 100 mM in DMSO. Using analogous methods described in the “High-content screening strategy” section, a 12-point dose response was constructed with concentrations ranging from < 100 nM to 100 µM of each compound, with a final working DMSO concentration of 0.1%. All compounds were tested using four replicates, and 12-positive (WT HeLa) and 12-negative (DMSO) controls were included on each screening plate to ensure assay robustness. MLF2-GFP was induced with 0.5 µg/mL of Dox 8 h before fixation, and compounds were added 3 h prior to fixation (5 h after initial Dox addition). Cells were washed, fixed, stained, and imaged as described above. All dose-response images were analyzed using an analogous *CellProfiler* + *R*-script pipeline.

The effect of each concentration was interpolated (*R*, *drc* & *ggplot*) using a four-parameter log-logistic function for each compound: 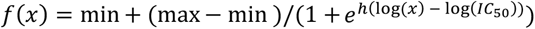, where x is the concentration, min represents the lower asymptote, max represents the higher asymptote, *IC*_50_ is the concentration at which response is 50% of the way between min and max, and *h* is the hill coefficient.^97,103^ The percent toxicity was calculated by fitting the additive inverse of the nuclear health function values to the same model.

### Pathway enrichment analysis

Gene Ontology (GO) term enrichment analysis was used for the top 100 CRISPR gene hits.^60^ The gene/chemical hits were separately analyzed in *R* to determine enriched biological process pathways using the enrichGO function (*clusterProfiler*).^104^ Enriched pathways were compared against an annotation database (org.Hs.eg.db). The top 30 pathways were then plotted using the pheatmap function and additionally plotted to an interactive .html graphic using heatmaply.^105^ Pathways were grouped by hierarchical clustering using Euclidean distance with complete linkage. To explore the interactive heatmap, please visit the <u>Schlieker Lab GitHub page</u> and download the corresponding .html files under the “Interactive Figures” folder.

### Immunofluorescence and confocal microscopy

HeLa cells were grown on 13 mm round coverslips and fixed with 4% (v/v) paraformaldehyde (Thermo Fisher Scientific), washed with 1x PBS (Gibco), permeabilized with 0.1% (v/v) Triton X-100 (Sigma-Aldrich) for 5 minutes, and blocked with a PBS mixture containing 2% (w/v) BSA and 2% (w/v) glycine. Primary antibodies were diluted in blocking buffer and added dropwise to coverslips for a one-hour incubation. After washing 3x with blocking buffer, fluorescent secondary antibodies diluted in blocking buffer were added for an additional hour, followed by a 1x PBS wash. Coverslips were then stained using a 1:2000 dilution of Hoechst 33342 (Life Technologies). Cells were then washed 3x with PBS before mounting onto slides (Fluoromount-G, Southern Biotech). Primary antibodies were diluted 1:500: anti-HA produced in rat (11867423001, Roche), anti-K48 produced in rabbit (ZRB2150, Millipore Sigma), and anti-Mab414 produced in mouse. Secondary antibodies were conjugated to Alexa Fluor 568 (Life Technologies).

Epifluorescence images were captured using a Zeiss AXIO Observer Z1 inverted fluorescence microscope using a ×63/1.40 oil objective. All confocal images were acquired at Yale’s Imaging Facility on Science Hill using a Zeiss LSM 880 laser scanning confocal microscope with a C Plan-Apochromat ×63/1.40 oil objective. Images were analyzed with *CondenScreen* and compiled for publication using Zeiss v3.4 and Image J (Fiji) software.

### Validation of hit compounds

Based on the dose-response results, we focused validation efforts on four compounds: PZ (IC_50_ = 601.6 nM), celastrol (IC_50_ = 124.8 nM), disulfiram (IC_50_ = 10.8 µM), and plumbagin (IC_50_ = 7.41 µM). To determine the optimal concentrations using a tighter set of concentrations, MLF2-GFP expression was induced in 4TorKO HeLa cells (0.5 µg/mL Dox, 8 h) and treated with either DMSO or one of four different concentrations/compounds: 2x below the IC_50_, IC_50_, 2x above the IC_50_, or 8x above the IC_50_. Compounds were added 3 h before fixation. For each condition, images (n > 50 cells/condition) were captured on a Zeiss AXIO Observer Z1 inverted fluorescence microscope and analyzed using our *CellProfiler* pipeline. *R Studio* was used to cluster MLF2-GFP foci into single cells grouped by treatment condition and construct jittered boxplots. To determine the effect of compounds on K48-ubiquitinylated proteins accumulation within condensates, we incubated cells with compounds for 3h (PZ, 3 µM; celastrol, 3 µM; plumbagin, 15 µM; and disulfiram, 50 µM), followed by immunofluorescent labeling with *α*-K48 primary antibody (1:500). Images were captured and analyzed as described above. High-resolution representative images were acquired using confocal microscopy (Zeiss LSM 880).

To determine differential protein expression, we constructed a 4TorKO HeLa cell line with a Dox inducible MLF2-GFP-P2A-tdTomato marker, where tdTomato serves as a diffuse secondary reporter for global protein synthesis. Cells were induced with 0.5 µg/mL Dox and treated with their respective compound for 6 h before fixation. Imaging was carried out on the Zeiss AXIO and the Zeiss LSM 880 for quantification and high-resolution, respectively. The integrated intensity of both GFP and tdTomato signal was quantified (n > 50) with *CellProfiler* and *R*. To ascertain the relative influence of global protein expression inhibition on the reduced number of MLF2-GFP foci, we derived the following empirical equation to determine the condensate normalization score 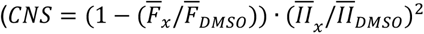, where 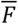 and 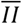 are the average number of NE condensate foci and the average integrated intensity of the secondary marker, respectively, for either the tested condition (*x*) or the DMSO control. The scoring system is designed to prioritize compounds that robustly reduce condensate foci without affecting tdTomato reporter expression and penalizes those that have little effect on foci or substantially suppress reporter expression. It penalizes modulators that have either little effect on foci or those that substantially suppress reporter expression. The weighting is particularly sensitive to changes in tdTomato intensity due to the squared exponent, thereby strongly penalizing compounds which impair protein expression. A low CNS-score suggests non-specific or off-target activities, such as decreased transcription, translation, or increased biomarker turnover.

### Guide RNA-mediated KO experiments

Each gene KO consisted of a pool of 3 distinct, synthetic sgRNAs, each targeting a different portion of the same gene (Horizon Discovery). Gene knock-out was conducted using either DharmaFECT or DharmaFECT Duo Transfection Reagent (Horizon discovery) using Tor1A KO + Tor1B KO as a positive control and NT sgRNAs as a negative control (Horizon discovery). The sgRNAs were transfected into stably-integrated Cas9 expressing HeLa cells and incubated for 72 h with a final sgRNA concentration of 25 nM. The cells were treated with 0.5 µg/mL Dox to induce the MLF2-GFP expression 6 h before fixation.

The pooled sgRNA sequences used were: ZNF335 sgRNAs (SG-016173, Horizon Discovery), _GATGATGGTGGCGCCGCCCA_, _GTGCCAGGGTGATCTGCGGT_, _ACCAAATGGACCAGGTGCCA_; RNF26 sgRNAs (SG-007060, Horizon Discovery), _TATCATGGAGGCAGTGTACC_, _GCATGACATCCTCTCGAAGA_, _CAGTGTACCTGGTAGTGAAT_; TOR1A sgRNAs (SG-011023, Horizon Discovery) _TATGAACATGGCTTTCTGGT_, _TATGAAGATGGACCTCGCAC_, _AGTGCAATGTGGCCACAAAC_; and TOR1B sgRNAs (SG-014203-01, Horizon Discovery), _AGTGCAGAGTCGATACAAAC_, _CAAGAGCCGTCTTAGTTATA_, _TTGGCGGGCCGGAAGAAAGA_.

### Sodium arsenite and 1,6-hexanediol treatment

To determine the co-localization of MLF2 into stress granules, Lipofectamine 2000 was used to co-transfect MLF2 with either G3BP1-GFP (Addgene #135997) + MLF2-HA, TDP-43-tdTomato-HA (Addgene #28205) + MLF2-GFP-Flag (Addgene #230986), or pcDNA3.1 empty vector control. Next day (16 h post-transfection), HeLa cells were treated with NaAsO_2_ (0.5 mM) and incubated for 30 min. Cells were fixed for 20 min with PFA, stained, and imaged as outlined. For hexanediol treatment, cells were incubated in complete DMEM medium containing 5% (v/v) 1,6-hexanediol (CAS 629-11-8) or 5% (v/v) 2,5-hexanediol (CAS 2935-44-6) for 5 min before fixation and immunofluorescence staining as described.

### Electron Microscopy

Electron microscopy was performed at the Center for Cellular and Molecular Imaging, Yale School of Medicine, using our previously described workflow (Laudermilch et al., 2016).^28^ Briefly, cells were fixed for 1 h in 2.5% glutaraldehyde in 0.1 M sodium cacodylate buffer, pH 7.4. Cells were rinsed and scraped in 1% gelatin and centrifuged in a 2% agar solution. Chilled cell blocks were processed with osmium and thiocarbohydrazide-osmium ligand as described (West et al., 2010), and samples were embedded in Durcupan ACM resin (Electron Microscopy Science). Polymerization was performed by incubating samples at 60°C overnight. These blocks were cut into 60 nm sections with a Leica UltraCut UC7 and stained with 2% uranyl acetate and lead citrate on Formvar/carbon-coated grids. Samples were visualized with an FEI Tecnai Biotwin TEM at 80 kV, and pictures were taken with Morada CCD and iTEM (Olympus) software.

### Statistical Analysis

Genetic and chemical hits were ranked based on their Z-Score or BZ-Score, respectively, as described in detail within the *Statistical analysis of screen data* section. GO pathways were ranked using a p-value cutoff of 0.05, after controlling for false discovery rate using the Benjamini-Hochberg (BH) procedure. The significance of all chemical and genetic validation experiments was assessed using a Bonferroni multiple comparisons corrected Mann–Whitney U test with > 50 cells per condition individually analyzed. Unless otherwise stated, asterisks indicate Bonferroni corrected p-values using a Mann-Whitney U test where *: p <= 0.05; **: p <= 0.01; ***: p <= 0.001; and ****: p <= 0.0001.

## Funding

This work was funded by the DOD PR200788 (C.S.) awarded by the U.S. Department of Defense, Department of the Army, along with support from the Dystonia Medical Research Foundation. Additionally, this work was supported by the NIH T32 GM145469 (D.P.), the European Molecular Biology Organization (EMBO) ALTF 910-2022 (E.F.F.K), and the Dutch Research Council (NWO) 019.222EN.007 (E.F.E.K.) This work utilized ImageXpress Micro 4 high content imager that was purchased with funding from a National Institutes of Health SIG grant 1S10OD032384. We thank Morven Graham and the Center for Cellular and Molecular Imaging (CCMI) Electron Microscopy Facility for the collection of TEM images and Shawn Ferguson for sharing stress granule marker reagents. The funders had no role in study design, data collection and analysis, decision to publish or preparation of the manuscript.

## Author contributions

Conceptualization: DP, CM, EFEK, YVS, CS

Methodology: DP, CM, SM, EFEK, VT, ND

Investigation: DP, CM, SM, EFEK, VT, ND, CS

Computational Pipeline: DP, VT

Machine Learning: DP

Data Analysis: DP

Supervision: CS

Writing—original draft: DP

Writing—review & editing: DP, CS, EFEK, CM, SM, ND, YVS

## Competing interests

Authors declare that they have no competing interests.

## Data and materials availability

All data are available in the main text or the supplementary materials. The computational *CondenScreen* pipeline and machine learning source code are available on the Schlieker Lab GitHub page:https://github.com/SchliekerLab/Condenscreen.git. At the time of publication, all raw screening images will be uploaded to the EMBL-EBI BioImage Archive: https://www.ebi.ac.uk/bioimage-archive/.

## Notes

https://github.com/SchliekerLab/Condenscreen.git

### Competing Interest Statement

The authors have declared no competing interest.

### Summary of Updates

Data files, figures, and additional experiments have been added since the initial submission.

